# Layer-Specific Vulnerability is a Mechanism of Topographic Map Aging

**DOI:** 10.1101/2022.05.29.493865

**Authors:** Alicia Northall, Juliane Doehler, Miriam Weber, Stefan Vielhaber, Stefanie Schreiber, Esther Kuehn

## Abstract

Topographic maps form a critical feature of cortical organization, yet are poorly described with respect to their microstructure in the living aging brain. We acquired quantitative structural and functional 7T-MRI data from younger and older adults to characterize layer-wise topographic maps of the primary motor cortex (M1). Using parcellation-inspired techniques, we show that qT1 and QSM values of the hand, face, and foot areas differ significantly, revealing microstructurally-distinct cortical fields in M1. We show that these fields are distinct in older adults, and that myelin borders between them do not degenerate. We further show that the output layer 5 of M1 shows a particular vulnerability to age-related increased iron, while layer 5 and the superficial layer show increased diamagnetic substance, likely reflecting calcifications. Taken together, we provide a novel 3D model of M1 microstructure, where body parts form distinct structural units, but layers show specific vulnerability towards increased iron and calcium in older adults. Our findings have implications for understanding sensorimotor organization and aging, in addition to topographic disease spread.

## 1. Introduction

Topographic maps are a hallmark feature of the human brain, covering nearly half of the cortical surface (Sereno et al., 2022). Cortical microcircuits of topographic maps, such as columns and layers, therefore reveal critical insights into the fundamental mechanisms of aging and neurodegeneration. While macrostructural changes in volume and thickness are often described in cortical aging (Thambisetty et al., 2010; Storsve et al., 2014), such measures show inconsistent relationships with behavior and their mechanistic origins are often unclear (Martinez et al., 2015). Microstructural features of topographic aging, reliably measured using quantitative imaging, provide more specific information that can be better linked to *ex-vivo* human data and animal evidence. Nevertheless, there has rarely been any systematic description of the cortical layer microcircuits of topographic maps in living older adults.

Cortical layers serve as functionally-relevant units, where deep layers typically control output functions and superficial layers typically integrate information from other regions, with distinct cytoarchitectural profiles that are often uniquely affected by pathology. For example, disrupted cross-laminar processing in layer 5 has been shown to precede cell death in aging mice (Lison et al., 2014). While such animal studies have long recognised the importance of layer-specific mechanisms in aging, such success in the specificity of *in-vivo* human aging has been limited by resolution constraints. Topographic map architectures are also relevant for human aging, where local and global changes in the functional map architecture influence functional integration and separation, and are linked to behavior (Liu et al., 2021). The importance of topographic maps also extends more generally, where functional and structural map maintenance has been related to phantom limb pain in amputees (Makin et al., 2013).

Recent advances in ultra-high field MRI at 7 Tesla (7T-MRI) provide the sub-millimetre resolution required to investigate cortical layers *in-vivo* (Kuehn & Pleger, 2020). 7T-MRI has successfully been used to parcellate the human sensorimotor cortex (Dinse et al., 2015), detect input and output signal flows in M1 (Huber et al., 2017), and detect layer-specific low-myelin borders between adjacent topographic areas in the sensorimotor cortex (Kuehn et al., 2017). While 7T-MRI is a suitable tool to describe the microstructural architecture of topographic maps *in-vivo*, it has not yet been applied to a group of healthy older adults. The mechanisms that underlie topographic map aging in humans are therefore yet to be elucidated.

We take the primary motor cortex (M1) as a model system to characterize *in-vivo* microstructural aging of topographic maps. M1 is the thickest (3-4mm) cortical region (McColgan et al., 2020), therefore providing greater relative resolution per layer compared to any other cortical region. In addition, M1 has a clear large-scale topographic organization and a high signal-to-noise ratio in MR images. The structural architecture of M1 is complex and demands a comprehensive consideration of local variations in microstructure. Although M1 is classically depicted as one homogeneous cortical field (Brodmann, 1909), there are microstructural gradients across two dimensions of the cortical surface (Glasser et al., 2016). Recent research suggests that M1 may be subdivided into separate cortical fields corresponding to topographic areas (Sereno et al., 2022). The microstructure of M1 also varies across a third dimension, cortical depth, according to the distinct cytoarchitecture of cortical layers (Kuehn & Sereno, 2018). M1 is characterized by a particularly thick layer 5 that contains the heavily-myelinated Betz cells that connect directly to the spinal cord to control motor output. Layer 5, which can be subdivided into layer 5a and layer 5b, is often implicated in neurodegeneration (McColgan et al., 2020) and may therefore be relevant in microstructural topographic map aging.

There are a number of open questions associated with the microstructure of M1 in younger and older adults. First, it is unclear whether the different topographic areas representing major body parts would have similar or distinct microstructural profiles. This is important to clarify before investigating how aging affects M1 microstructure. Evidence from mice research has demonstrated varying effects of aging across topographic areas, where for example representations of the hindpaw are more vulnerable to age-related dedifferentiation than those of the forepaw (David-Jürgens et al., 2008). The inhomogeneity of M1 microstructure is further complicated by the presence of low-myelin borders that divide major topographic areas, such as the hand and face (Glasser et al., 2016; Kuehn et al., 2017). The role of these borders in cortical aging, particularly in the enlargement of body part representations, is currently unknown. Finally, it also needs to be clarified which cortical layers, and which topographic areas, are affected most in older adults to understand the precise architecture of topographic map aging.

To characterize microstructural aging with respect to layers and topographic areas, we applied parcellation-inspired techniques to sub-millimetre 7T-MRI data in healthy younger and older adults. We employed validated in-vivo proxies of cortical myelin (quantitative T1 (qT1), Stüber et al., 2014), iron (positive QSM (pQSM)), Langkammer et al., 2012) in addition to a proxy of diamagnetic substance (negative QSM (nQSM)), as measures of cortical microstructure. Together with functional localisers, we extracted microstructural profiles of the major cortical fields of M1 (lower limb (LL), upper limb (UL), face (F)) (termed according to Glasser et al., 2017). Age-related differences in pQSM and nQSM values were of particular interest, since previous studies have shown increased iron (Acosta-Cabronero et al., 2016; Betts et al., 2016) and calcium (Jang et al., 2021; Kim et al., 2022) in aging. Increased iron has been related to poorer motor function in healthy older adults (Sullivan et al., 2009), whereas the biological source of the nQSM signal is currently under debate (see more details in the discussion). In order to investigate whether such age-effects are uniformly present across cortical layers or are layer-specific, we estimated anatomically-relevant cortical compartments *in-vivo*. Our approach was based on a comparison between *in-vivo* and *ex-vivo* M1 data (Huber et al., 2017), therefore providing a reasonable approximation of anatomically-relevant compartments and their computations (Persichetti et al., 2020). However, please note that there is a conceptual difference between our definition and the definition based on *ex-vivo* data, where cytoarchitecture is considered (Brodmann, 1909; Vogt & Vogt, 1919).

We applied a systematic approach to characterize microstructure within the different cortical fields of M1 representing the LL, UL, and F areas in a sample of healthy adults. (1) We first hypothesized that M1, in healthy younger adults, is comprised of microstructurally-distinct cortical fields corresponding to topographic areas (as suggested by (Flechsig, 1920), also see (Glasser et al., 2016; Kuehn et al., 2017; Sereno et al., 2022)). (2) We further hypothesized that the microstructure of these cortical fields is also distinct in older adults, but that (3) the low-myelin borders between them (previously shown in younger adults, see Kuehn et al., 2017) are degenerated in older adults. In addition, (4) we hypothesized that iron and diamagnetic substance in older adults would be elevated in a layer- and topographic area-specific way given the above cited evidence in rodents. (5) Finally, we hypothesized that those changes would show a relation to body part-specific motor function.

For all analyses, the focus is on the left (dominant) hemisphere, since all participants were right-handed. However, we also tested whether effects replicate for the right (non-dominant) hemisphere, which is particularly relevant for diseases in which pathology can onset in either hemisphere, or is related to handedness (Turner et al., 2011). To the best of our knowledge, this is the first study to apply recently-introduced techniques of *in-vivo* 3D parcellation (Alkemade et al., 2022; Kuehn & Sereno., 2018) to group brain data of different age groups. This will allow us to uncover the fundamental aspects of microstructural M1 architecture and aging in the human brain.

## 2. Materials and methods

### 2.1. Participants

40 healthy volunteers including 20 younger adults (< 35 years of age; 8 female) and 20 older adults (> 70 years of age; 11 female) were enrolled in the present study. After data quality check (see section 2.5.1 for details), a total of 35 participants, including 17 younger adults (8 female) with a mean age of 25 years (*SD* = 2.7 years) and 18 older adults (11 female) with a mean age of 71 years (*SD* = 4.0 years) remained for analysis. Additional demographic information and group differences are shown in **Table 1**. Participants were recruited from the DZNE database in Magdeburg and were paid seven euros per hour for the behavioral tests, and 30 euros for each MRI session. All participants were right-handed and exclusion criteria included chronic illnesses, neurological medications and contraindications to 7T-MRI (e.g. tattoos, metallic implants, tinnitus). We ensured that participants had intact sensorimotor function, had no chronic pain condition or other disorders that would affect their tactile or motor behavior. The study was approved by the local Ethics Committee and all participants gave written informed consent.

**Table 1.** Demographic information and group differences for younger and older adults involved in the myelin analysis. Independent-samples t-tests were used to investigate group differences. Differences marked * and ** indicate significance at the .05 and .001 p-value levels, respectively. MoCA: Montreal Cognitive Assessment.

|  | Younger adults (n = 17) | Older adults (n = 18) | Group difference |  |  |
| --- | --- | --- | --- | --- | --- |
|  | Mean (SD) | Mean (SD) | *df* | *t* | *Sig.* |
| Age (years) | 24.65 (2.69) | 71.06 (4.04) | 34 | -39.77 | 10<sup>-5**</sup> |
| Education (years) | 17.24 (2.31) | 16.06 (2.26) | 34 | 1.53 | .136 |
| MoCA | 28.88 (1.11) | 27.41 (2.21) | 33 | 2.45 | .020* |

### 2.2. Procedure

Data acquisition took place between August 2016 and November 2021, and included two MRI scanning sessions and one behavioral session: (1) Structural MRI session, (2) Functional MRI session, (3) Motor behavior session.

### 2.3. MRI Data Acquisition

Structural MRI data were acquired with a 7T-MRI scanner equipped with a 32-channel head coil (MAGNETOM, Siemens Healthcare), located in Magdeburg, Germany. We collected two MP2RAGE sequences (Magnetisation Prepared 2 Rapid Gradient Echoes) (Marques et al., 2010), including a MP2RAGE slab image (covering M1 and S1) with a 0.5 mm isotropic resolution (208 transversal slices, repetition time = 4800 ms, echo time = 2.62 ms, inversion time TI1/TI2 = 900/2750 ms, field-of-view read = 224 mm, bandwidth = 250 Hz/Px, GRAPPA 2, flip angle = 5◦/3◦, phase oversampling = 0%, slice oversampling = 7.7%) and a whole-brain MP2RAGE image with an isotropic resolution of 0.7 mm (sagittal slices, repetition time = 4800 ms, echo time = 2.01 ms, inversion time TI1/TI2 = 900/2750 ms, field-of view read = 224 mm, bandwidth = 250 Hz/Px, GRAPPA 2, flip angle = 5◦/3◦). We also collected a whole-brain susceptibility-weighted (SWI) image with a 0.5 mm isotropic resolution (transversal slices, repetition time = 22 ms, echo time = 9 ms, field-of view read = 192 mm, bandwidth = 160 Hz/Px, GRAPPA 2, flip angle = 5◦/3◦). The total scan time for the structural MRI session (session 1) was approximately 60 minutes.

During the functional MRI session (session 2), functional and additional SWI data were collected. The same sequence and parameters as above were used to collect SWI slab images (covering M1 and S1) with a 0.5 mm isotropic resolution. In addition, we obtained whole-brain functional images with a 1.5 mm isotropic resolution (81 slices, field-of-view read = 212 mm, echo time = 25 ms, repetition time = 2000 ms, GRAPPA 2, interleaved acquisition) using an EPI gradient-echo sequence.

### 2.4. Experimental Design

The functional imaging (session 2) involved a blocked-design paradigm, where the participants were instructed to move their left or right foot, left or right hand, or tongue. Participants were trained on the movements outside the scanner and wore fingerless braces, covering the hands and forearms, to reduce large movement of the hands during the precise movement of the fingers and other body parts. Instructions were shown on a screen inside the scanner (gray background, black color). They were instructed to prepare for movement (e.g. ‘prepare right hand’) before carrying out the movement (e.g. ‘move right hand’) for 12 seconds, followed by 15 seconds of rest. Each movement was repeated four times, resulting in a total of 20 trials. The total scan time for the MRI session (2) was approximately 90 minutes.

### 2.5. Image Processing

#### 2.5.1. Data Quality Inspection

The MP2RAGE data of n = 5 adults (n = 3 younger, n = 2 older) were excluded due to low SNR or severe truncation artifacts affecting M1, leaving a total of n = 35 participants (n = 17 younger, n = 18 older) for the myelin analyses. In addition, the SWI data of n = 5 adults (n = 3 younger, n = 2 older) were excluded due to severe motion and truncation artifacts affecting M1, leaving a total of 31 participants (n = 14 younger, n = 16 older) for the SWI analyses.

#### 2.5.2. Structural Preprocessing

Structural images were processed using CBS Tools (version 3.0.8) (Bazin et al., 2014) implemented in MIPAV (version 7.3.0) (McAuliffe et al., 2001). We first registered the slab MP2RAGE image to the whole-brain MP2RAGE image using the ‘Optimized Automated Registration’ module in MIPAV and ANTs (version 1.9.x) (Avants et al., 2011). Registration quality in the sensorimotor areas were checked by two raters before proceeding. The slab and whole-brain images were then merged, resulting in whole-brain MP2RAGE images with improved resolution in the slab region. Using CBS Tools, the MP2RAGE skull stripping module was employed to remove extra-cranial tissue, and the MP2RAGE dura estimation module was used to estimate the dura mater, which was manually refined in ITK-SNAP. The topology-preserving anatomical segmentation (TOADS) algorithm (Bazin & Pham, 2008) was used to segment the UNI images into different tissue types based on a voxel-by-voxel probability approach. The cortical reconstruction, using implicit surface evolution (CRUISE) module, was then used to estimate the boundaries between the WM and GM and between the GM and CSF (Han et al., 2004), resulting in levelset images. Using these images, we created the subject-specific cortical surfaces used for mapping the microstructural measures.

#### 2.5.3. SWI Processing

Quantitative Susceptibility Maps (QSM) were reconstructed using QSMbox (version 2.0) (Acosta-Cabronero et al., 2018). We chose not to normalize the QSM values, since previous studies have demonstrated comparable aging effect sizes between normalized and non-normalized data, both with 3T- (Acosta-Cabronero et al., 2016) and 7T-MRI (Betts et al., 2016). The reconstructed QSM images were registered to the merged qT1 images using the automated registration tool in ITK-SNAP (version 3.8.0), with manual refinement (prioritizing alignment in M1) where necessary. Registration quality between the QSM and MP2RAGE images was checked by two raters before proceeding. We separated the QSM data into positive QSM (pQSM) and negative QSM (nQSM) values following a previously described approach (Betts et al., 2016).

#### 2.5.4. M1 Masks

ROI masks of M1 in both hemispheres were manually delineated, based on the MP2RAGE image, using the manual segmentation tool in ITK-SNAP. Based on anatomical landmarks, the M1 region was defined to include all relevant topographic areas (foot, hand, face/bulbar). The omega-shaped knob, known as the ‘hand knob’, serves as a reliable and robust landmark of the hand area in M1 with 97-100% accuracy (Yousry et al., 1997). We first identified the hand knob (located at the anterior wall of the central sulcus) in axial slices and masked all slices in which it was visible. Then, to include the face (bulbar) area, we masked slices inferior to the hand knob until the precentral gyrus was no longer visible (Donatelli et al., 2019). Finally, to include the foot area, we masked slices superior to the hand knob until the precentral gyrus was no longer visible. Since the foot area extends superiorly to the longitudinal fissure (Penfield & Rasmussen, 1950), we masked the paracentral lobule (PCL) while avoiding the supplementary motor area and the primary somatosensory cortex (Spasojević et al., 2013). Finally, the masks were refined in the coronal and sagittal views.

#### 2.5.5. Surface Mapping

We divided the cortex into 21 layers according to the equivolume approach implemented in the volumetric layering module (Waehnert et al., 2014; 2016) of CBS Tools, using the volume-preserving layering method and an outward layering direction. We then sampled qT1, acquired using the MP2RAGE sequence, and signed QSM values at each layer, before mapping the values onto inflated cortical surfaces (see **Fig. 1**) using the surface mesh mapping module with the closest-point method.

**Figure 1.**
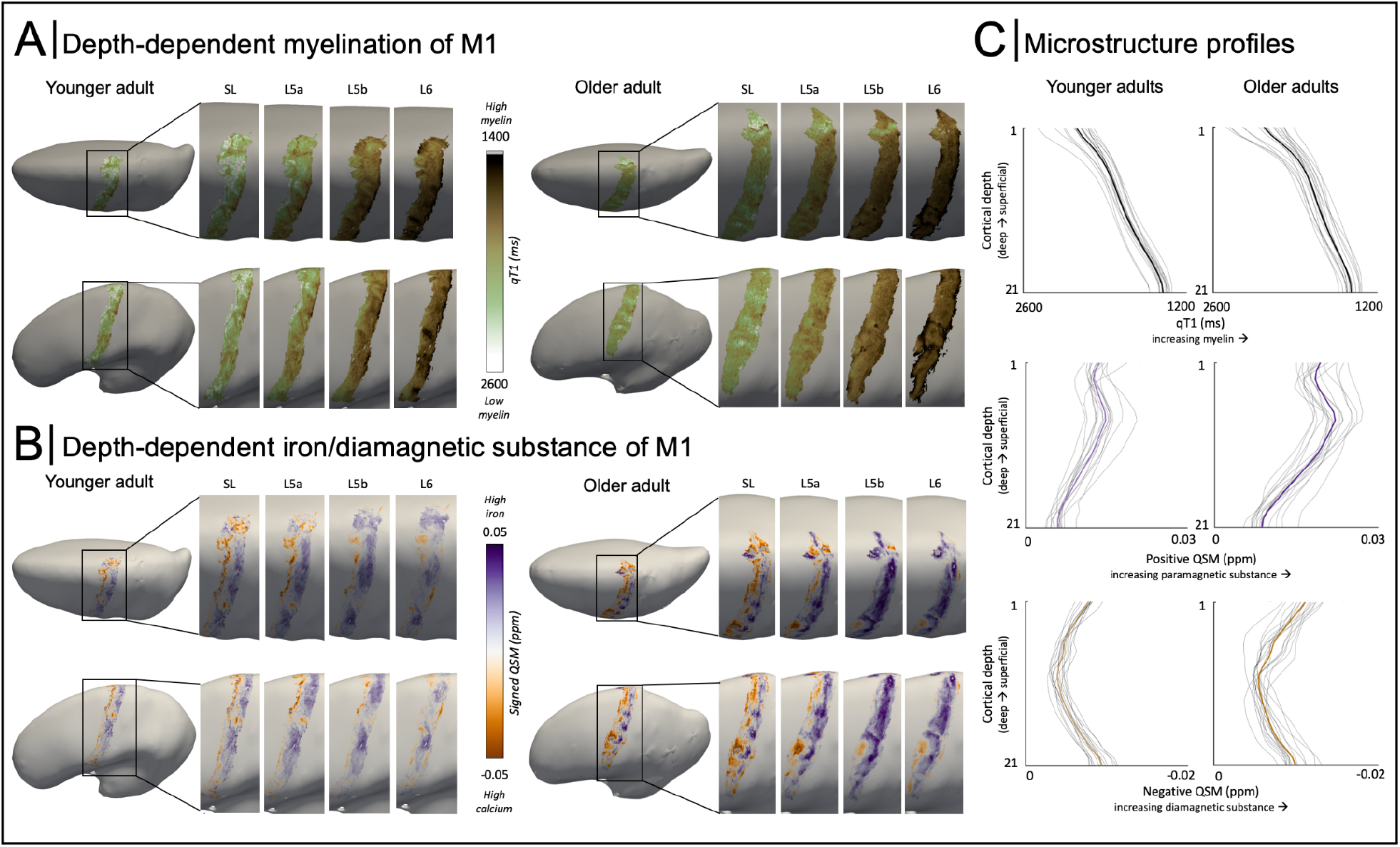
3D Microstructural Profiles of Left Primary Motor Cortex (M1). (A) Mapped qT1 values in M1 at each of the four anatomically-relevant cortical compartments (‘layers’) (Ls - superficial layer, L5a - layer 5a, L5b - layer 5b, L6-layer 6) for one example younger adult and one example older adult. (B) Mapped signed QSM values in M1 at each of the four anatomically-relevant cortical compartments (‘layers’), for one example younger adult and one example older adult. Note that positive QSM values represent iron, whereas negative values reflect diamagnetic substance/calcium (C) ‘Raw qT1’ (i.e., not ‘decurved qT1’) profiles, positive QSM profiles, and negative QSM profiles for younger and older adults mapped at 21 cortical depths; the group mean is shown in bold color whereas individual data are shown in gray.

#### 2.5.6. Defining Cortical Layers

We calculated mean curvature using the simple curvature function in ParaView (version 5.8.0) and regressed this out of qT1 values for each layer, on an individual basis, following a previously published approach (Sereno et al., 2013). Using group-averaged decurved qT1 profiles, we applied a data-driven approach to define anatomically-relevant cortical compartments (Huber et al., 2017) (see Fig. 2). Layer 5 (L5) was identified based on a plateau in decurved qT1 after a steep increase in decurved qT1, reflecting the sharp increase in myelin content from the superficial layers to the heavily-myelinated L5 (depths 4-13). In addition, we distinguished between layer 5a (L5a) (4-7) and layer 5b (L5b) (8-13) based on the presence of two small qT1 dips that are considered to represent the two layer compartments (Huber et al., 2017). All depths above L5a were then labeled as the superficial layer (Ls, 1-3), which we suggest includes anatomical layers 3 and 4 but not anatomical layers 1 and 2, since the latter are particularly sparse and inaccessible with MRI (Huber et al., 2017). Finally, we defined layer 6 (L6, 14-17) based on a sharp decrease in decurved qT1, after which values plateaued again indicating the presence of WM (18-21). Taken together, we here refer to Ls, L5a, L5b and L6 when referring to cortical compartments that presumably refer to anatomical layers 3 and 4 (Ls), anatomical layer 5a (L5a), anatomical layer 5b (L5b) and anatomical layer 6 (L6), where, however, an exact delineation of the layers would need *ex-vivo* validation.

**Figure 2.**
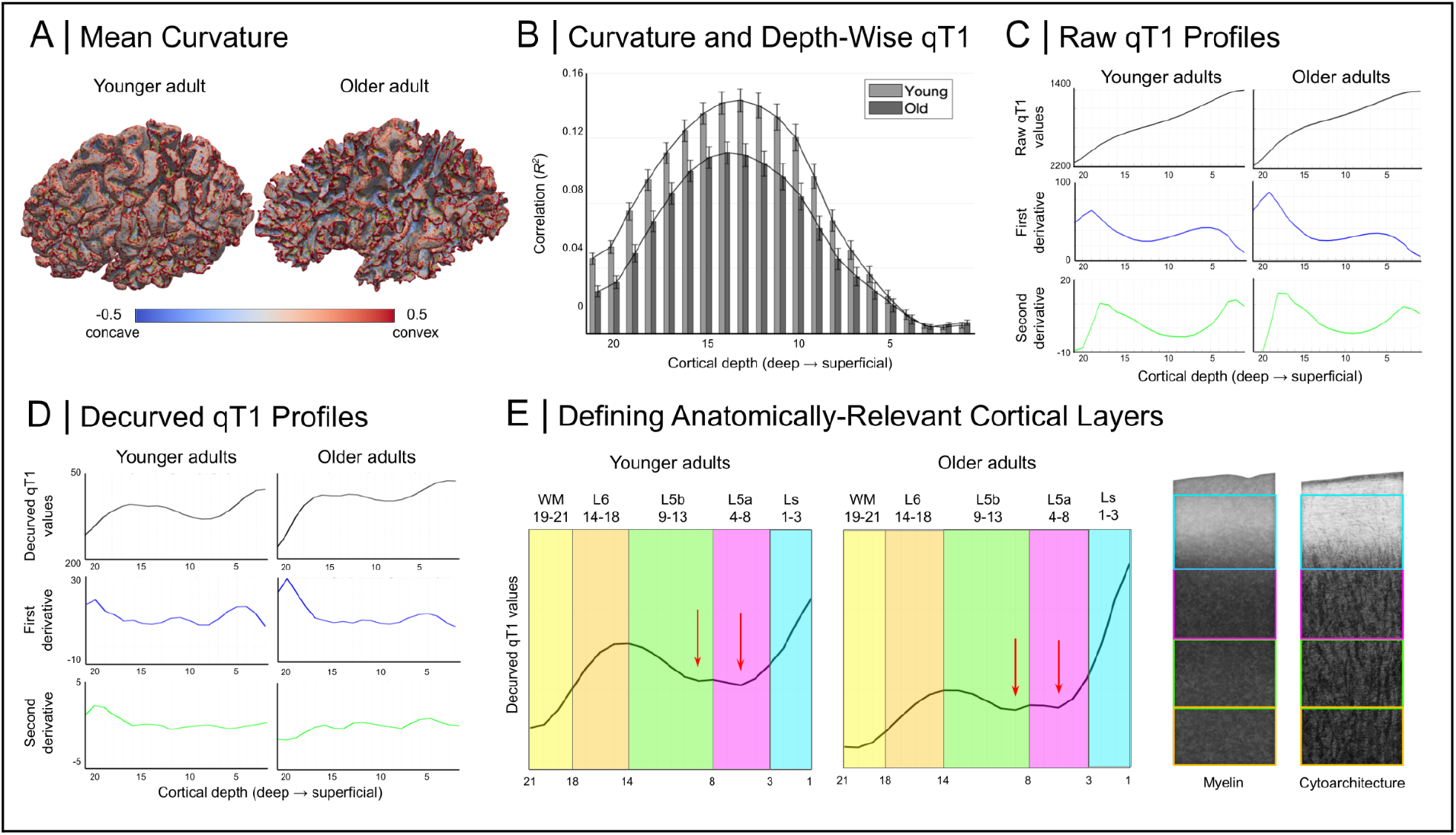
Identifying Anatomically-Relevant Cortical Compartments in Left Primary Motor Cortex (M1). (A) Mapped mean cortical curvature for one example younger adult and one example older adult. (B) Cortical depth-wise correlations between qT1 values and mean curvature for younger adults (light gray) and older adults (dark gray). Note that the 21 correlations shown correspond to the initially-extracted 21 depths. (C) Group mean ‘raw qT1’ profiles along with the first and second derivatives. (D) Group mean ‘decurved qT1’ profiles along with the first and second derivatives. (E) According to a previously published approach (Huber et al., 2017), we identified four anatomically-relevant compartments (‘layers’) based on ‘decurved qT1’ (Ls - superficial layer; L5a - layer 5a; L5b - layer 5b; L6 - layer 6). L5a and L5b were distinguished based on the presence of two small qT1 dips at the plateau of ‘decurved qT1’ values (indicating L5), while L6 was identified based on a sharp decrease in values before a further plateau indicating the presence of white matter. We show our layer approximations over schematic depictions of M1 myelin (left) (from Dinse et al., 2015) and cell histological staining (right) (Vogt & Vogt, 1919). Note that these cortical layers are defined based on in-vivo MRI data and may not correspond exactly to the anatomical layers as defined by ex-vivo myelo- and cytoarchitecture. Also note that here, low qT1 values represent high myelin whereas in the other graphs in the article, values are plotted reversed such that high values represent high myelin.

#### 2.5.7. Functional Data Processing

The functional data were retrospectively motion-corrected using the Siemens ‘MoCo’ correction. The data were preprocessed using SPM12, including smoothing with a Gaussian kernel of 2 mm, slice-timing to correct for differences in image acquisition time between slices, and realignment to reduce motion-related artifacts. The functional volumes were averaged and manually registered to the qT1 images using ITK-SNAP, based on anatomical landmarks. First-level analysis was used to generate t-statistic maps (t-maps) based on contrast estimates for each body part (e.g. left hand = [1 0 0 0 0]). The peak cluster of each t-map was saved as a binary mask. In the few cases where the peak cluster only included the face area of one hemisphere (tongue movement typically elicits a single peak cluster including the face area of both hemispheres), we included the largest cluster in the face area of the other hemisphere in the resulting mask. The t-maps and peak cluster masks were then registered to the qT1 images in ANTs, using the registration matrices previously generated in ITK-SNAP, before being mapped onto the same inflated cortical surfaces as used for the structural data (see **Fig. 1**). We applied the binarised peak cluster masks to the t-maps to create functional localisers which contained t-values only in the peak cluster area. We refined the localisers by removing overlapping voxels between different localisers, where the body part with the highest t-value retained the overlapping voxel (i.e., ‘winner-takes-it-all’ approach). These refined functional localisers (shown in **Fig. 3A**) were then binarised and multiplied with the layer-wise qT1 and QSM values, resulting in values for each cortical field at four different cortical layers.

**Figure 3.**
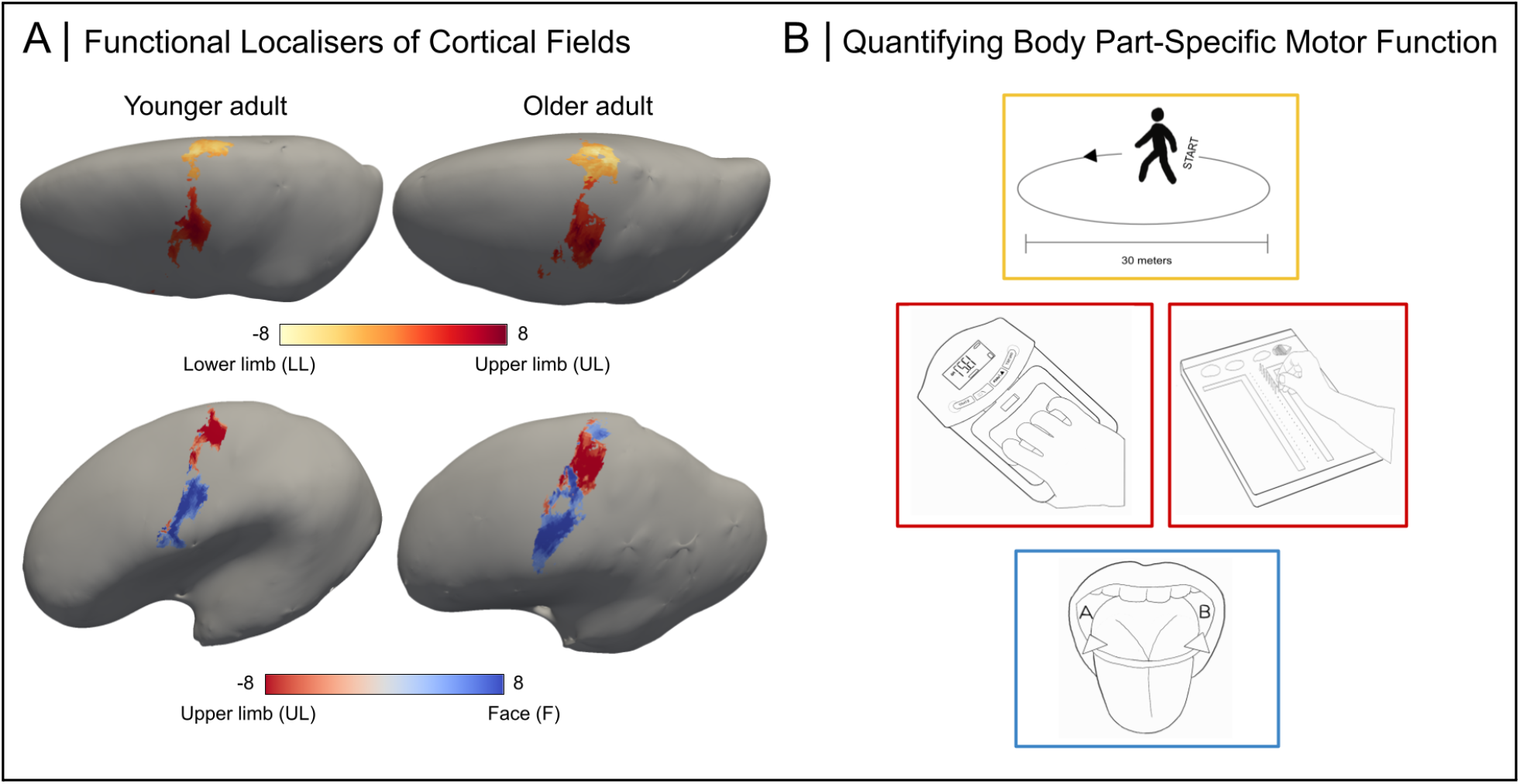
Body Part-Specific Behavioral Motor Impairments in Older Adults. (A) Functional localisers of large-scale body part representations in M1 for one example younger adult and one example older adult. (B) Example behavioral tests to quantify body part-specific motor function (Lower limb shown in yellow color: 6MWT; Upper limb shown in red color: Hand dynamometer Purdue pegboard; Face shown in blue color: tongue movement task). Note that the O’Connor pegboard is not visualized here.

#### 2.5.8. Myelin Border Analysis

Based on previous evidence of low-myelin borders between the hand and face areas of M1 (Glasser et al., 2016; Kuehn et al., 2017), we defined the myelin border based on the highest qT1 value (lowest myelin) between the locations of the peak t-values of the UL and F localisers. We extracted the qT1 values at the border and calculated the average body part qT1 values (UL+F/2).

### 2.6. Behavioral Tests of Motor Function

A subset (n = 27) of participants (12 younger, 15 older) underwent body part-specific behavioral tests of motor function (see **Fig. 3B**). The 6-Minute-Walking-Test (6MWT) was used to measure walking distance, as an estimate of gross motor function of the legs (Enright, 2003). Participants were instructed to walk as far as possible around a 30-meter measuring tape in six minutes, without speed-walking or running. The total distance walked was multiplied by weight in kilograms as previously described (Bernstein et al., 1994). We used a hand-held dynamometer to measure hand strength twice in each hand, before averaging to result in one measurement per hand (Peolsson et al., 2001). We used pegboards to measure hand and finger dexterity, more specifically the Purdue pegboard test (Lafayette Instrument, model 32020A; Tiffin & Asher, 1948) and the O’Connor pegboard test (Lafayette Instrument, model 32021; Fleishman, 1953). An automated open-source tongue tracking tool (Tongue Tracker, TT), was used to quantify kinematic features of the tongue based on short video clips of lateral tongue movement, as an estimate of bulbar motor function (Northall et al., 2022, code: https://github.com/BudhaTronix/Automated-Video-Analysis-Tool-for-Quantifying-Bulbar-Function). Participants were instructed to open their mouth and move their tongue between their mouth commissures as fast as possible, while being video recorded for a minimum of five seconds of correct movement. We used TT to automatically extract the average duration (‘TT speed’) of tongue movements and the number of errors (‘TT errors’) made by the participant.

### 2.7. Statistical Analysis

Statistical analyses were performed using IBM SPSS Statistics (version 26, IBM, USA). Mixed-effects ANOVAs were used to assess between-group (age group) and within-group (layer, cortical field) differences for each measure of cortical microstructure (qT1, pQSM, nQSM). The significance level for all statistical tests was set to the 5% threshold (p < .05). Significant main effects and interactions of the ANOVAs were investigated using post-hoc tests and the Holm-Bonferroni method (Holm, 1979) was used to correct for multiple comparisons. In the cases where the assumption of sphericity was violated (Sig. < 0.05) according to Mauchly’s test, we applied a sphericity correction. The correction used depended on the epsilon value calculated by the Greenhouse-Geisser correction (Field, 2013). If ε < 0.75, then the Greenhouse-Geisser correction was used, while the Huyn-Feld correction was used if ε > 0.75. As a result of this correction, the reported degrees of freedom and F-values are sometimes not reported as whole numbers. Two-tailed independent-samples t-tests were used to measure group differences in the size of the myelin border and in behavioral measures of motor function, while one-tailed independent-samples t-tests were used to measure group differences in the mean dice coefficient of the overlap between functional representations. Linear regression models were used to investigate the influence of demographic variables (gender, education, cognitive function) on the cortical microstructure of M1.

## 3. Results

### 3.1. Body Part-Specific Behavioral Motor Impairments in Older Adults

We quantified body part-specific motor function in younger and older adults (see section 2.6 for details and **Fig. 3B** for visualization, see Supplementary Table 1 for statistics).

### 3.2. Extracting 3D Microstructural Profiles of M1 with High Precision

#### 3.2.1. Identifying Anatomically-Relevant Cortical Compartments in M1

We described the topographic microstructure of M1 using qT1 as a proxy for cortical myelin content (Stüber et al., 2014), pQSM as a marker for cortical iron content (Langkammer et al., 2012), and nQSM as a measure of cortical diamagnetic substance (Deh et al., 2018). A sub-millimeter isotropic resolution of 0.5 mm allowed us to characterize intracortical contrast with high precision (see **Fig. 1** for an overview of extracted data). Given known correlations between curvature and layer-wise qT1 (Sereno et al., 2013) (see **Fig. 2A & 2B** for our data), we followed a previous approach to regress out effects of curvature from ‘raw qT1’ values (see **Fig. 2C)** for each layer, resulting in ‘decurved qT1’ (see **Fig. 2D**, see Supplementary Fig. 1 for the right hemisphere). We then applied a data-driven approach (Huber et al., 2017) to divide M1 into four anatomically-relevant compartments, based on intracortical myelin content, that we refer to as ‘layers’ (Ls = superficial layer; L5a = layer 5a; L5b = layer 5b; L6 = layer 6; see **Fig. 2E**).

#### 3.2.2. Demographic Variables Do Not Predict Variance in M1 Microstructure

To investigate if demographic variables (gender, education, cognitive function) are associated with the microstructure of M1, we performed regression analyses on each microstructural measure (qT1, pQSM, nQSM) averaged across cortical fields (dependent variable), with demographic variables (gender, years of education, MoCA score) as predictors. We show that none of the demographic variables significantly predict qT1 values (*R*^2^ = .10, *F*(3, 33) = 1.05, *p* = .385; gender (*t* = -.21, *p* = .833), education (*t* = -1.01, *p* = .321), MoCA (*t* = 1.26, *p* = .219)), pQSM values (*R*^2^ = .23, *F*(3, 28) = 2.54, *p* = .079; gender (*t* = -1.32, *p* = .199); education (*t* = .86, *p* = .401; MoCA ((*t* = -2.63, *p* = .140)) or nQSM values (*R*^2^ = .08, *F*(3, 28) = .75, *p* = .531); gender (*t* = .97, *p* = .340), education (*t* = -.74, *p* = .468), MoCA (*t* = 1.30, *p* = .205)).

### 3.3. M1 is Comprised of Distinct Cortical Fields

In order to target our first hypothesis (In healthy younger adults, M1 is comprised of microstructurally-distinct cortical fields corresponding to topographic areas), we first tested whether M1 is best described as a single cortical field (as suggested by Brodmann, (Brodmann, 1909)), or whether it is comprised of several fields (as suggested by Flechsig, 1920; see also Sereno et al., 2022). It has previously been reported that the UL and F areas of M1 are separated by low-myelin borders (Glasser et al., 2016; Kuehn et al., 2017). However, it has not yet been investigated whether the microstructure of these areas significantly differs, which would be necessary to define them as separate ‘cortical fields’. To investigate this, we computed ANOVAs with the within-subjects factors cortical field (LL, UL, F) and layer (Ls, L5a, L5b, L6) on qT1, pQSM and nQSM values as *in-vivo* proxies of cortical microstructure in younger adults.

With respect to qT1 values, there are significant main effects of layer (*F* _(1.17, 18.63)_ = 576.42, *P* < 10^−16^, η_p_^2^ = .97) and cortical field (*F*_(1.30, 20.83)_ = 5.34, *P* = .024, η _p_^2^ = .25), as well as a significant interaction between layer and cortical field (*F* _(2.12, 33.98)_ = 12.78, *P* < 10^−5^, η_p_ ^2^ = .44; note that we report values of the dominant (left) hemisphere in the main text, but see Supplementary Tables 2 & 3 for the corresponding analyses on the non-dominant (right) hemisphere). The main effect of layer was expected and is driven by a significant decrease in qT1 (i.e., an increase in cortical myelin) with cortical depth that is due to the high myelination in the deep cortex near the white matter (Dinse et al., 2015). More specifically, Ls shows higher qT1 (i.e., less myelin) than L5a (Ls = 2083.17 ± 115.92; L5a = 1898.49 ± 72.21; *df* = 16, *t* = 14.60, *P* < 10^−10^, *d* = 3.54, 95% CI [2.23 4.84]), L5a shows higher qT1 than L5b (L5b = 1743.68 ± 54.60; *df* = 16, *t* = 21.59, *P* < 10^−13^, *d* = 5.24, [3.37 7.09]), and L5b shows higher qT1 than L6 (L6 = 1569.05 ± 61.75; *df* = 16, *t* = 36.41, *P* < 10^−17^, *d* = 8.83, [5.76 11.89]).

The main effect of cortical field is due to significantly lower qT1 in the LL area compared to the UL area (LL = 1818.24 ± 73.04; UL = 1824.57 ± 72.48; *df* = 16, *t* = -3.87, *P* = .001, *d* = -.94, 95% CI [-1.50 -.35]). The significant interaction between layer and cortical field is driven by the LL area showing significantly lower qT1 than the UL area in L5a (LL = 1894.08 ± 71.82; UL = 1902.77 ± 72.26; *df* = 16, *t* = -5.41, *P <* 10^−5^, *d* = -1.31, [-1.96 -.65]) and L5b (LL = 1739.48 ± 54.64; UL = 1746.50 ± 54.63; *df* = 16, *t* = -4.87, *P <* 10^−4^, *d* = -1.18, [-1.80 -.55]). In addition, the LL area shows significantly lower qT1 than the F area in L6 (LL = 1560.76 ± 63.27; F = 1582.22 ± 60.91; *df* = 16, *t* = -6.03, *P <* 10^−5^, *d* = -1.46, 95% CI [-2.14 -.76]) and the UL area shows significantly lower qT1 than the F area in L6 (UL = 1564.16 ± 62.64; *df* = 16, *t* = -4.22, *P* = .001, *d* = -1.02, [-1.61 -.42]).

With respect to pQSM values, there are significant main effects of layer (*F*_(1.20, 15.65)_ = 184.05, *P <* 10^−4^, η_p_^2^ = .93) and cortical field (*F*_(2, 26)_ = 10.35, *P <* 10^−10^, η_p_^2^ = .44), as well as a significant interaction between layer and cortical field (*F*_(1.91, 24.88)_ = 6.32, *P <* .007, η_p_^2^ = .33, see Supplementary Tables 4 & 5 for the right hemisphere). Note that higher pQSM reflects more iron (Langkammer et al., 2012). The main effect of layer is driven by Ls showing higher pQSM (i.e., more iron) than L5a (Ls = .0139; L5a = .0117, *df* = 13, t = 10.33, P < 10^−7^, *d* = 2.76, [1.58 3.92]), L5a showing significantly higher pQSM than L5b (L5b = .0073, *df* = 13, *t* = 13.91, *P <* 10^−9^, *d* = 3.72, [2.20 5.22]), and L5b showing significantly higher pQSM than L6 (L6 = .0051, *df* = 13, *t* = 9.39, *P <* 10^−7^, *d* = 2.51, 95% CI [1.14 3.59]).

The main effect of cortical field is due to higher pQSM in the F area compared to the UL area (F = .0100; UL = .0093; *df* = 13, *t* = 3.53, *P* = .004, *d* = .94, 95% CI [.30 1.57]) and significantly higher pQSM in the F area compared to the LL area (LL = .0092; *df* = 13, *t* = 4.80, *P <* 10^−4^, *d* = 1.28, [.55 1.99]). The significant interaction between layer and cortical field is driven by the F field showing higher pQSM values than the LL field and the UL field in Ls (F = .0146 ; LL = .0137; *df* = 13, *t* = 4.04, *P* = .001, *d* = 1.08, [.40 1.73]; UL = .0133; *df* = 13, *t* = 4.41, *P* = .001, *d* = 1.17, [.48 1.86]) and L5a (F = .0124 ; LL = .0114; *df* = 13, *t* = 5.30, *P <* 10^−4^, *d* = 1.42, 95% CI [.65 2.16]; UL = .0113; *df* = 13, *t* = 4.34, *P* = .001, *d* = 1.16, [.46 1.83]).

With respect to nQSM values, there are significant main effects of layer (*F*_(1.41, 18.35)_ = 235.32, *P <* 10^−13^, η _p_^2^ = .95) and cortical field (*F*_(2.26,_ = 5.05, *P* = .014, η _p_^2^ = .28), but there is no significant interaction between layer and cortical field (*F*_(2.5, 32.50)_ = 1.50, *P* = .235, η _p_^2^ = .10, see Supplementary Tables 6 & 7 for the right hemisphere). Note that lower (i.e. more negative) nQSM reflects more diamagnetic tissue contrast (Deh et al., 2018). The main effect of layer is driven by L5a showing significantly higher nQSM (i.e., less diamagnetic tissue) than L5b (L5a = -.0055 ± .0008, L5b = -.0088 ± .0013, *df* = 13, *t* = 19.55, *P <* 10^−11^, *d* = 5.22, [3.17 7.27]) and L5b showing significantly higher nQSM than L6 (L6 = -.0130 ± .0016, *df* = 13, *t* = 20.81, *P <* 10^−11^, *d* = 5.56, 95% CI [3.38 7.73]), while Ls does not show significantly different nQSM to L5a (Ls = -.0060 ± .0008; *df* = 13, *t* = -1.78, *P* = .098, *d* = -.48, [-1.02 .09]). The main effect of cortical field is due to lower nQSM in the F area compared to the LL area (F = -.0086 ± .0008; LL = -.0081 ± .0011; *df* = 13, *t* = -2.58, *P* = .023, *d* = -.69, 95% CI [-1.27 .09]) and the UL area (UL = -.0083 ± .0010; *df* = 13, *t* = -2.17, *P* = .049, *d* = -.58, [-1.14 .002]), although these differences were not significant at the Holm-Bonferroni-corrected significance levels.

Taken together, we show that the F, UL and LL areas of M1 show significant differences in all three quantitative tissue contrasts in younger adults, which we summarize in a novel 3D model of the microstructural architecture of healthy M1 (see **Fig. 4A**). We show that the F area, overall, is characterized by high iron and diamagnetic substances. We also show layer-specific differences between cortical fields, for example, the F area shows low myelin in L6. In line with our hypothesis (1), our results indicate systematic differences in the microstructural profiles of the cortical fields of M1 in younger adults; we therefore refer to these areas as ‘cortical fields’ in the following text when investigating age-related changes in the M1 architecture (i.e., LL field, UL field, F field), in line with a recently-introduced parcellation atlas based on functional data (Sereno et al., 2022).

**Figure 4.**
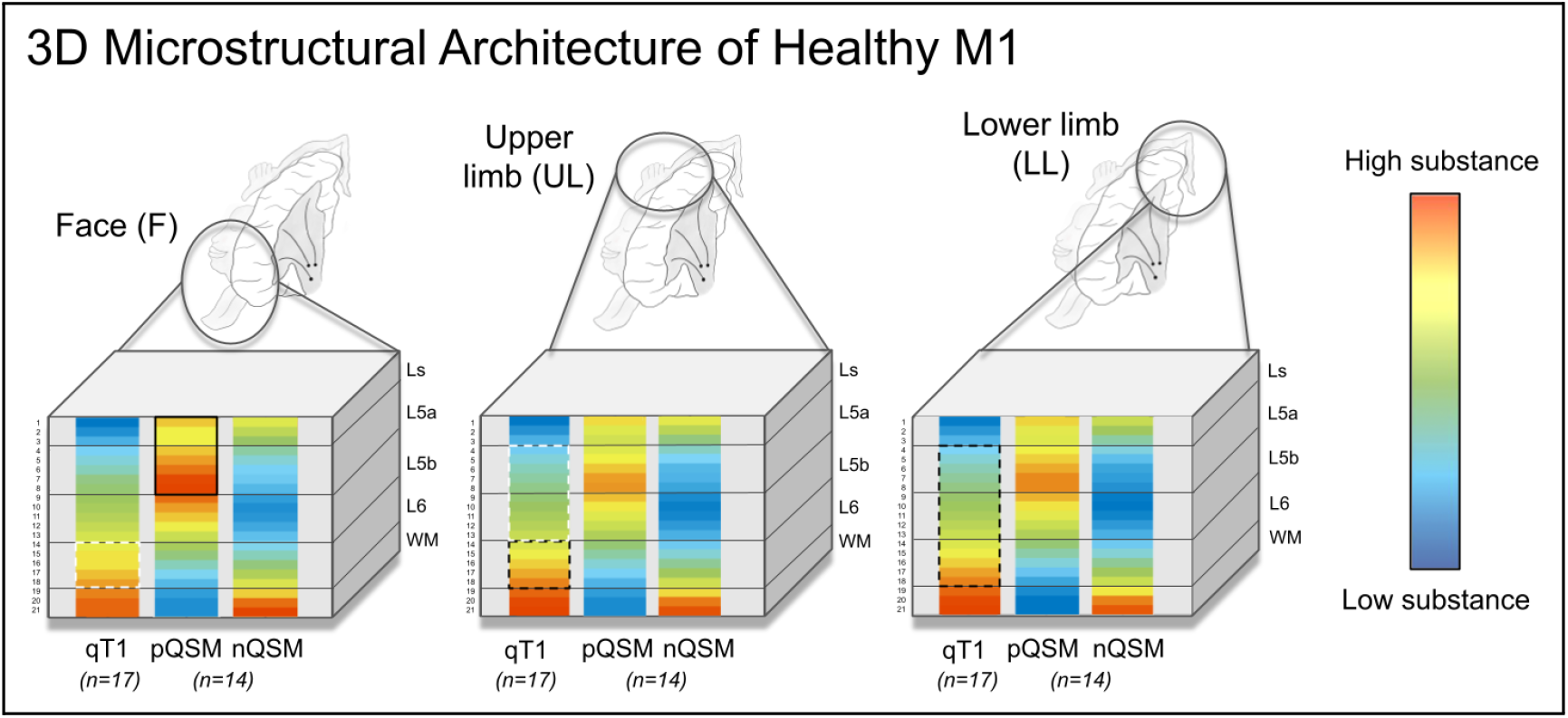
3D Microstructural Architecture of the Healthy Left Primary Motor Cortex (M1). Cubes show the raw data across all 21 cortical depths, to highlight which microstructural features characterize the face (F), upper limb (UL) and lower limb (LL) fields of the dominant (left) hemisphere for younger adults only. Black solid and white solid lines indicate that the highlighted layer contains the highest and lowest substance, respectively, compared to the corresponding layer of both other cortical fields; black dotted and white dotted lines indicate that the highlighted layer contains lower substance, respectively, compared to the corresponding layer of one other cortical field (see section 3.3 for statistics).

### 3.4. Cortical Fields Are Distinct in Older Adults

To target our second hypothesis (The microstructure of cortical fields is distinct in older adults), we calculated ANOVAs on qT1 and nQSM values with the factors cortical field (LL, UL, F), layer (Ls, L5a, L5b, L6) and age (younger, older). In short, we replicate the differences between cortical fields across age groups (see details below, see **Fig. 5**), and, critically, we do not find a significant interaction between age and cortical field or between age, layer and cortical field on qT1 values (interaction between age and cortical field: *F*_(2, 33)_ = 2.22, *P* = .117, η_p_^2^ = .06; interaction between age, layer, and cortical field: *F*_(6, 198)_ = 1.18, *P* = .320, η_p_^2^ = .03) or nQSM values (interaction between age and cortical field: *F*_(2, 28)_ = .60, *P* = .550, η_p_^2^ = .02; interaction between age, layer, and cortical field: *F*_(6, 168)_ = .20, *P* = .977, η_p_^2^ = .01). This implies that the microstructural differences in qT1 and nQSM values between the F, UL and LL areas, as reported above, do not show significant age-related changes. Note that also for the right hemisphere, no such interactions were found (see Supplementary Tables 8 & 10). Also note that the pQSM (iron) results are reported below.

**Figure 5.**
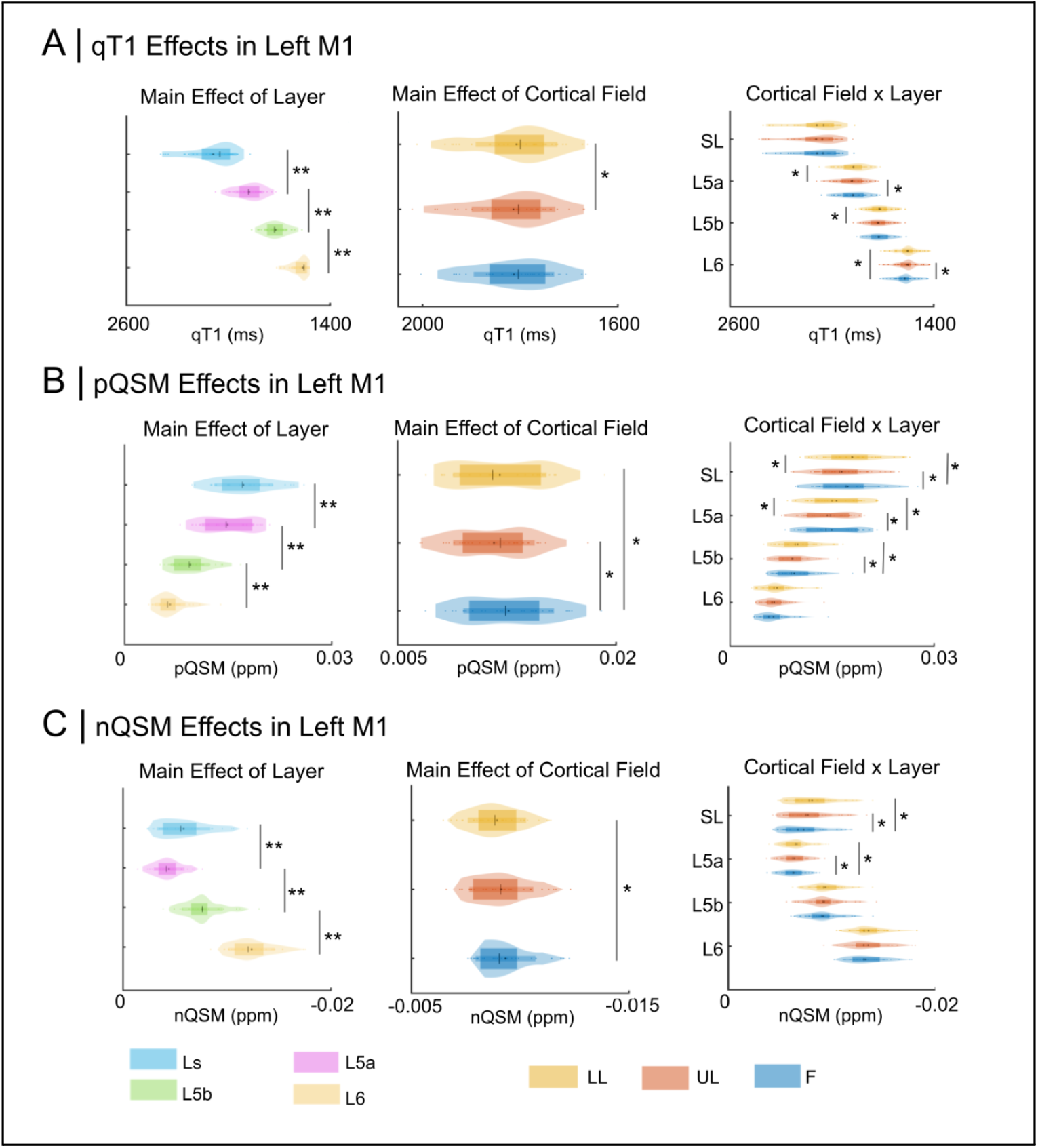
Microstructurally Distinct Cortical Fields in Left Primary Motor Cortex (M1). Layer- and cortical field-specific effects on (A) qT1 values as a proxy for cortical myelin content (Stuber et al., 2014), (B) pQSM values as a proxy of cortical iron content (Langkammer et al., 2012), and (C) nQSM values as a measure of diamagnetic tissue contrast (Deh et al., 2018). Main effects: * and ** indicate significance at the 5% and 1% significance levels, respectively. Interactions: * indicates statistical significance at 5% level, corrected for multiple comparisons using the Holm-Bonferroni method.

More specifically, an ANOVA with the factors cortical field (UL, LL, F), layer (Ls, 5a, 5b, 6) and age (younger, older) on qT1 values revealed significant main effects of layer (*F*_(1.18, 38.90)_ = 781.22, *P <* 10^−28^, η_p_^2^ = .96) and cortical field (*F*_(2, 66)_ = 3.67, *P* = .031, η_p_^2^ = .10), as well as a significant interaction between layer and cortical field (*F*_(2.34, 77.04)_ = 15.75, *P* < 10^−7^, η_p_^2^ = .32, see **Fig. 5A** and see Supplementary Tables 8 & 9 and Supplementary Fig. 2A for the right hemisphere). As expected, the main effect of layer is driven by a significant decrease in qT1 with cortical depth, where Ls shows higher qT1 than L5a (Ls = 2088.64 ± 138.93; L5a = 1876.59 ± 83.43; *df* = 34, *t* = 17.02, *P <* 10^−18^, *d* = 2.88, 95% CI [2.12 3.63]), L5a shows higher qT1 than L5b (L5b = 1720.50 ± 67.50; *df* = 34, *t* = 30.14, *P <* 10^−26^, *d* = 5.10, [3.84 6.34]), and L5b shows higher qT1 than L6 (L6 = 1559.86 ± 65.59; *df* = 34, *t* = 34.81, *P <* 10^−28^, *d* = 5.89, [4.45 7.31]).

The main effect of cortical field is driven by significantly lower qT1 in the LL field compared to the UL field (LL = 1807.69 ± 83.83; UL = 1813.76 ± 85.63; *df* = 34, *t* = -3.18, *P* = .003, *d* = -.54, 95% CI [-.89 -.18]). The significant interaction between layer and cortical field is driven by the LL field showing significantly lower qT1 than the UL field in L5a (LL = 1873.29 ± 82.67; UL = 1882.57 ± 85.57; *df* = 34, *t* = -4.50, *P <* 10^−5^, *d* = -.76, 95% CI [-1.13 -.38]) and L5b (LL = 1716.51 ± 68.02; UL = 1724.45 ± 68.12; *df* = 34, *t* = -5.06, *P <* 10^−3^, *d* = -.86, [-1.24 -.46]), and the LL field showing significantly lower qT1 than the F field in L6 (LL = 1553.19 ± 66.60; UL = 1570.33 ± 65.46; *df* = 34, *t* = -5.85, *P <* 10^−6^, *d* = -.99, [-1.39 -.58]) across age groups. In addition, the F field shows significantly lower qT1 than the UL field in L5a (F = 1873.90 ± 83.39; UL = 1882.57 ± 85.57; *df* = 34, *t* = -3.09, *P* = .004, *d* = -.52, 95% CI [-.87 -.17]), while the opposite effect is shown in L6, where the UL field shows significantly lower than the F field (UL = 1556.08 ± 66.30; F = 1570.33 ± 65.46; *df* = 34, *t* = -5.76, *P <* 10^−6^, *d* = -.57, [-1.37 .57]). Note that the results of the ANOVA on pQSM values (see **Fig. 5B**) are reported in **Tables 2 & 3**, since we describe the age-related pQSM effects in more detail in section 3.6 (see Supplementary Tables 13-15, and Figure 2B for the right hemisphere).

**Table 2.** Main effects of layer and cortical field, and the interaction between layer and cortical field on pQSM values in younger (n = 14) and older (n = 16) adults. See section 3.6 for main effects of age and significant interactions with age. * and ** indicate statistical significance at the 5% and 1% significance levels, respectively.

| Factor | df | Sum of squares | Mean square | F | p | $\eta_p^2$ |
| --- | --- | --- | --- | --- | --- | --- |
| Layer | 1.31 | .006 | .002 | 299.99 | $10^{-21**}$ | .92 |
| cortical field | 2 | $10^{-5}$ | $10^{-5}$ | 22.17 | $10^{-8**}$ | .44 |
| Layer x cortical field | 2.44 | $10^{-5}$ | $10^{-6}$ | 9.80 | $10^{-5**}$ | .26 |

**Table 3.** Post-hoc t-tests for significant (non-age) ANOVA effects on pQSM values in younger (n = 14) and older (n = 16) adults. (Ls = superficial layer; L5a = layer 5a; L5b = layer 5b; L6 = layer 6) * indicates statistical significance at 5% level, corrected for multiple comparisons using the Holm-Bonferroni method.

| Pair | Paired differences |  |  |  |  |  |
| --- | --- | --- | --- | --- | --- | --- |
|  | df | Mean difference | Standard deviation | t | Sig. | d, [95% CI] |
| Layer; Cortical field |  |  |  |  |  |  |
| Ls-L5a | 29 | -.0029 | .0012 | -14.76 | 10<sup>-15*</sup> | -2.70 [-3.47 -1.91] |
| L5a-L5b | 29 | -.0054 | .0018 | -16.87 | 10<sup>-16*</sup> | -3.08 [-3.94 -2.21] |
| L5b-L6 | 29 | -.0023 | .0017 | -7.44 | 10<sup>-8*</sup> | -1.36 [-1.85 -.85] |
| F-UL | 29 | .0010 | .0009 | 5.88 | 10<sup>-6*</sup> | 1.07 [.62 1.52] |
| UL-LL | 29 | -.0004 | .0009 | -2.39 | .024 | -.44 [-.81 .06] |
| LL-F | 29 | -.0006 | .0007 | -4.75 | 10<sup>-5*</sup> | -.87 [-1.28 -.44] |
| Layer * cortical field |  |  |  |  |  |  |
| Ls: F-UL | 29 | .0016 | .0013 | 6.75 | 10<sup>-7*</sup> | 1.23 [.75 1.70] |
| Ls: UL-LL | 29 | -.0009 | .0010 | -4.86 | 10<sup>-5*</sup> | -.89 [-1.31 -.46] |
| Ls: LL-F | 29 | -.0007 | .0011 | -3.53 | .001* | -.65 [-1.04 -.25] |
| L5a: F-UL | 29 | .0014 | .0012 | 6.12 | 10<sup>-6*</sup> | 1.12 [.65 1.53] |
| L5a: UL-LL | 29 | -.0006 | .0011 | 3.13 | .004* | .57 [.18 .96] |
| L5a: LL-F | 29 | -.0007 | .0009 | -4.22 | 10<sup>-4*</sup> | -.77 [-1.18 -.36] |
| L5b: F-UL | 29 | .0007 | .0008 | 4.67 | 10<sup>-5*</sup> | .86 [.43 1.27] |
| L5b: UL-LL | 29 | -.0001 | .0011 | -1.14 | .264 | -.21 [-.57 .16] |
| L5b: LL-F | 29 | -.0005 | .0009 | -3.06 | .005* | -.56 [-.94 -.17] |
| L6: F-UL | 29 | .0004 | .0010 | 2.21 | .035 | .40 [.03 .77] |
| L6: UL-LL | 29 | .0001 | .0011 | .67 | .506 | .12 [-.24 .48] |
| L6: LL-F | 29 | -.0005 | .0011 | -2.73 | .011 | -.50 [-.88 -.12] |

The same ANOVA on nQSM values shows significant main effects of layer (*F*_(1.19, 33.41)_ = 169.24 and *P <* 10^−15^, η_p_^2^ = .86) and cortical field (*F*_(2, 56)_ = 9.42 and *P <* 10^−4^, η_p_^2^ = .25), as well as a significant interaction between layer and cortical field (*F*_(2.45, 68.45)_ = 3.60 and *P* = .025, η_p_^2^ = .11, see **Fig. 5C** and see Supplementary Table 10 & 11 and Supplementary Fig. 2C for right hemisphere). The main effect of layer reflects a U-shaped curve in nQSM with cortical depth, where Ls shows lower nQSM than L5a (Ls = -.0076 ± .0022, L5a = -.0064 ± .0013, *df* = 29, *t* = -3.99, *P <* 10^−4^, *d* = -.73, [-1.13 .32]), L5a shows higher nQSM than L5b (L5b = -.0092 ± .0017, *df* = 29, *t* = 12.58, *P <* 10^−13^, *d* = 2.30, [1.60 2.98]) and L5b shows higher nQSM than L6 (L6 = -.0133 ± .0019, *df* = 29, *t* = 35.59, *P <* 10^−25^, *d* = 6.50, 95% CI [4.79 8.20]).

The main effect of cortical field is driven by the F field showing significantly lower nQSM than the LL field (LL = -.0089 ± .0013; *df* = 29, *t* = -3.78, *P* = .001, *d* = -.69, [-1.08 -.29]). The significant interaction between layer and cortical field on nQSM values reveals that this effect is confined to Ls and L5a. More specifically, the F field shows significantly lower nQSM than the LL field and UL field in Ls (F = -.0080 ± .0024; LL = -.0072 ± .0022; *df* = 29, *t* = -3.88, *P* = .001, *d* = -.71, [-1.11 -.30]); UL = -.0076 ± .0023; *df* = 29, *t* = -3.86, *P* = .001, *d* = -.71, 95% CI [-1.10 -.30]). The F field also shows significantly lower nQSM than the LL field and UL field in L5a (F = -.0066 ± .0013; LL = -.0063 ± .0012; *df* = 29, *t* = -3.40, *P* = .002, *d* = -.62, [-1.01 -.23]); UL = -.0063 ± .0013; *df* = 29, *t* = -3.22, *P* = .003, *d* = -.59, 95% CI [-.97 -.20]).

In regards to age effects, there is a main effect of age on nQSM values (*F*_(1,28)_ = 12.88, *P* = .001, η_p_^2^ = .32, see **Fig. 6B**), which is driven by significantly lower values (i.e., more diamagnetic tissue contrast) in older adults compared to younger adults across all cortical fields and layers in M1 (younger adults = -.0083 ± .0001, older adults = -.0098 ± .0013, *df* = 28, *t* = -3.59, *P* = .001, *g* = -1.28, 95% CI [-2.04 -.50]. There is also a significant interaction between layer and age (*F*_(1.19, 28)_ = 6.19, *P* = .014, η _p_^2^ = .18, see **Fig. 6D**), revealing that older adults show lower nQSM values compared to younger adults specifically in Ls (older adults = -.0091 ± .0021; younger adults = -.0060 ± .0001; *df* = 28, *t* = -5.57, *P* < 10^−5^, *g* = -2.18, [-2.73 -1.02]) and L5a (older adults = -.0072 ± .0011; younger adults = -.0055 ± .0084; *df* = 28, *t* = -4.61, *P* < 10^−5^, *g* = -1.72, 95% CI [-2.45 -.81]). Note that we do not find a significant main effect of age, or a significant interaction between age and cortical layer, on nQSM values for the right hemisphere (see Supplementary Table 10).

**Figure 6.**
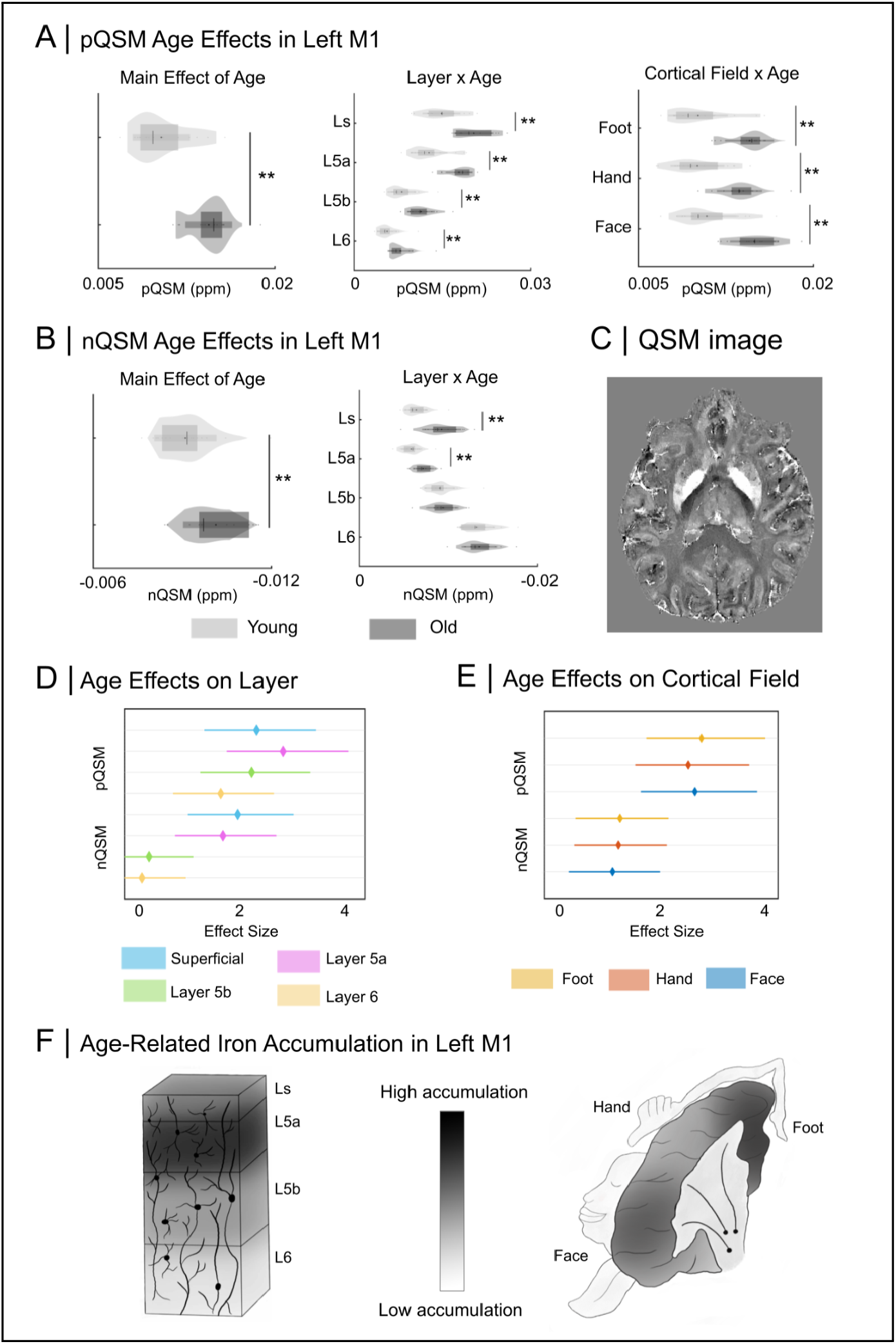
Age-Related Layer-Specific Iron and Diamagnetic Substance Increases in Left Primary Motor Cortex (M1). (A) Significant main effect of age, and significant interactions between layer and age, and between age and cortical field on pQSM values. (B) Significant main effect of age and significant interaction between age and layer on nQSM values. (C) Example whole-brain QSM image for a younger adult, showing high values (indicating high iron) in the basal ganglia. (D) Effect sizes for the age effects on layer QSM values, while (E) shows effect sizes for the age effects on cortical field QSM values. (F) Schematic diagram showing different vulnerability to iron increase in M1 across the cortical surface and with depth (Ls = superficial layer; L5a = layer 5a; L5b = layer 5b; L6 = layer 6). Main effects: * and ** indicate significance at the 5% and 1% significance levels, respectively. Interactions: * indicates statistical significance at 5% level, corrected for multiple comparisons using the Holm-Bonferroni method.

Overall, we show that the microstructural differences between the cortical fields (LL, UL, F) of M1, as reported above, can also be detected in older adults. In addition, older adults show more diamagnetic substance (nQSM) than younger adults. This effect is atopographic (i.e., even across cortical fields) but layer-specific (i.e., specific to Ls and L5a, see **Fig. 6B, 6D & 6E**).

### 3.5. Hand-Face Myelin Borders Is Not Degenerated in Older Adults

To test our third hypothesis (The low-myelin border between the hand and the face is degenerated in older adults), we compared averaged qT1 sampled in the UL and F fields with the averaged qT1 sampled at the UL-F border, at each layer, between younger and older adults. There are no significant age differences in any layer, but there is a trend in Ls towards a greater border in older adults (Ls: older adults = -778.14 ± 212.61; younger adults = -643.95 ± 216.08, *df* = 33, *t* = 1.85, *P* = .073, *g* = .63, 95% CI [-.06 1.30]; L5a: older adults = -589.47 ± 298.48; younger adults = -474.42 ± 216.16, *df* = 33, *t* = 1.30, *P* = .203, *g* = .43, [-.23 1.08]; L5b: older adults = -361.78 ± 171.87; younger adults = -363.58 ± 163.46, *df* = 33, *t* = -.03, *P* = .975, *g* = -.01, [-.66 .64]; L6: older adults = -307.50 ± 152.34; younger adults = -308.16 ± 120.93, *df* = 33, *t* = -.01, *P* = .989, *g* = -.01, [-.65 .64]; see **Fig. 7A**) (see Supplementary Table 12 and Supplementary Fig. 3 for the right hemisphere).

**Figure 7.**
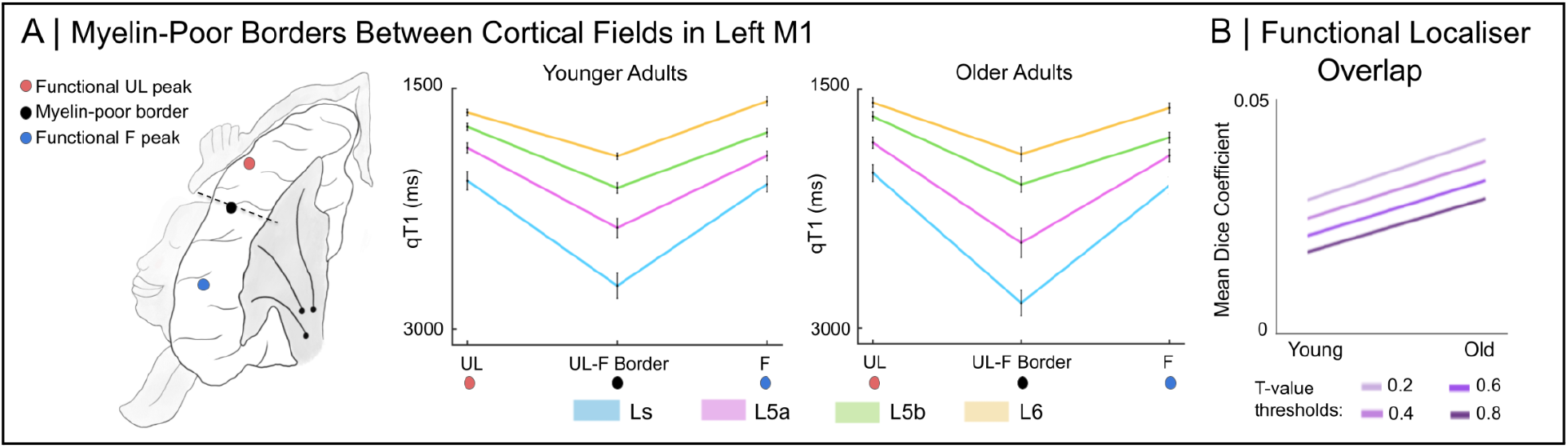
Stable Hand-Face Myelin Borders in Older Adults in Left Primary Motor Cortex (M1). (A) Layer-wise low-myelin (high qT1 value) border between the cortical fields of the upper limb (UL) and face (F) for both younger and older adults. In each group, we compare the mean body part qT1 values (hand+face/2) with the qT1 values at the hand-face border across layers. (B) Mean dice coefficients of the overlap between F and UL areas for younger and older adults, using different t-value thresholds to restrict the size of the functional representations.

To additionally investigate whether greater functional overlap between body part representations occurred in older adults, despite intact (or even slightly elevated) structural borders, we calculated the mean dice coefficient of the overlap between the functional localisers of the UL field and the F field (Dice, 1945). We compared the dice coefficients between younger and older adults, where the representations where thresholded at different t-values (threshold = .02: *df* = 27.12, *t* = -1.83, *P* = .079, *g* = -.59, 95% CI [-1.25 .07]; threshold = .04: *df* = 26.70, *t* = -1.77, *P* = .088, *g* = -.58, [-1.23 .09]; threshold = .06: *df* = 26.38, *t* = -1.77, *P* = .088, *g* = -.58, [-1.24 .09]; threshold = .08: *df* = 26.28, *t* = -1.78, *P* = .087, *g* = -.58, [-1.24 .09]; see **Fig. 7B**). While we found no significant differences, we show statistical trends (*P* < .1) at all thresholds, where older adults show greater overlap compared to younger adults. In addition, we show that the degree of this overlap is not significantly correlated with the size of the low-myelin UF-F border (averaged across layers) in older adults (n = 18), at any of the t-value thresholds (threshold = .02: *r* = -.27, *p* = .274; threshold = .04: *r* = -.21, *p* = .404; threshold = .06: *r* = -.27, *p* = .286; threshold = .08: *r* = -.19, *p* = .442).

Taken together, we do not confirm our hypothesis that a degenerated hand-face structural border in older adults would relate to larger cortical representations / to more overlap between the UL and F areas. We instead show a trend towards a more pronounced hand-face boundary in older compared to younger adults in the superficial layer of M1, whereas the other layers do not show a trend towards a difference. Please note that this effect in the superficial layer does not survive a correction for multiple comparisons.

### 3.6. Layer-specific Iron Accumulation in Older Adults’ M1

#### 3.6.1. Layer-Specific Vulnerability in Older Adults

To test our fourth hypothesis (Iron in older adults is elevated in a layer- and topographic area-specific way), we investigated whether increased iron in older adults’ M1 is uniformly present across cortical layers, or whether it occurs in particular cortical layers only. In addition, we investigated whether the age-related increase in iron is even across cortical fields, or whether some cortical fields show a selective vulnerability. We performed an ANOVA with the factors cortical field (LL, UL, F), layer (Ls, L5a, L5b, L6) and age (younger, older) on pQSM values, which served as a validated *in-vivo* proxy for cortical iron content (Langkammer et al., 2012).

As expected, there is a significant main effect of age on pQSM values (*F*_(1,28)_ = 56.24, *P <* 10^−8^, η_p_^2^ = .67), reflecting significantly higher pQSM values (i.e., more iron) in older adults compared to younger adults across M1 (older adults = .0143 ± .0019; younger adults = .0095 ± .0016, *df* = 28, *t* = 7.50, *P <* 10^−8^, *g* = 2.67, 95% CI [1.67 3.64], see **Fig. 6A** and see Supplementary Table 13 and Supplementary Fig. 4 for the right hemisphere). Additionally, there is also a significant interaction between layer and age on pQSM values (*F*_(1.31, 28)_ = 8.88, *P <* .003, η_p_^2^ = .24), which is driven by older adults showing the strongest effect size (largest Hedge’s *g*), that is, most iron accumulation in L5a (older = .0178 ± .0023; younger = .0117 ± .0023; *df* = 28, *t* = 7.42, *P* < 10^−8^, *g* = 2.72, 95% CI [1.65 3.61], see **Fig. 6A & 6D**). There is also a significant interaction between age and cortical field on pQSM values (*F*_(2, 28)_ = 3.89, *P <* .026, η_p_^2^ = .12), revealing that older adults show the strongest effect size, that is highest iron increase compared to younger adults, in the LL field (F field: older = .0149 ± .0020; younger = .0100 ± .0018; *df* = 28, *t* = 7.24, *P* < 10^−8^, *g* = 2.66, 95% CI [1.60 3.53]; UL field: older = .0137 ± .0018; younger = .0093 ± .0017; *df* = 28, *t* = 6.92, *P* < 10^−7^, *g* = 2.54, [1.50 3.40]; LL field: older = .0144 ± .0021; younger = .0092 ± .0016; *df* = 28, *t* = 7.56, *P* < 10^−8^, g = 2.81, [1.69 3.67]) (see **Fig. 6A & 6E**; see Supplementary Tables 14 and 15 for post-hoc tests for the right hemisphere). We have visualized the uneven age-related iron accumulation across the cortical layers and cortical fields of M1 in **Fig. 6F**. For the additional results of the ANOVA, see **Tables 2 & 3**.

#### 3.6.2. Age-Related Iron Differences are Independent of Cortical Atrophy

To investigate whether cortical atrophy can explain the age-related differences in pQSM values (iron content), we calculated the ANOVA on pQSM values, as described above, while controlling for mask size (i.e. cortical field mask sizes used to extract microstructural profiles). We show that the main effect of age remains significant (F (1,28) = 20.71, P < 10-5, ηp2 = .43). The interaction between age and cortical layer (F (1.31,28) = 3.66, P = .016, ηp2 = .12) also remains significant, and is still driven by older adults showing the strongest effect size (largest Hedge’s g), that is, most iron accumulation in L5a (older = .0014 ± .0029; younger = -.0024 ± .0024; df = 28, t = 4.60, P < 10-5, g = 1.64, 95% CI [.81 2.44]). Our findings are supported by previous evidence that age-related QSM effects are largely independent of brain atrophy measured with 7T-MRI (Betts et al., 2016).

Taken together, we show that increased iron in older adults’ M1, a hallmark of cortical aging (Hallgren & Sourander, 1958), is particularly pronounced in L5a, suggesting output layer vulnerability. We further show that this effect is independent of atrophy in the topographic areas of M1. We also show that other than the increased diamagnetic substance, age-related iron increase is not atopographic but topographic in the left hemisphere, because there was most age-related iron accumulation in the LL cortical field. This effect was not present in the right (non-dominant) hemisphere.

### 3.7. The Functional Relevance of M1 Microstructure

To target our fifth and final hypothesis (Microstructural M1 changes show a relation to body part-specific motor function), we performed correlational analyses for each cortical field, for younger and older adults separately, between microstructural measures and motor function (see **Fig. 8**). Given the subset of participants with behavioral data was small (n = 12 for younger adults, n = 15 for older adults) and the number of tests was large (n = 27), we report correlation coefficients without p-values to provide an overview of potential relationships. To ensure that higher values indicate ‘better performance’ or ‘more substance’ in all cases, Purdue pegboard scores (speed and pin drops), O’Connor pegboard scores (pin drops), TT scores (duration and errors) and qT1 values were reversed, while nQSM values were taken as absolute values. In this way, red fields in the matrix always indicate that ‘better performance’ relates to ‘more substance’ whereas blue values in the matrix always indicate that ‘better performance’ relates to ‘less substance’; this allows for an easier interpretation of the matrix. Of note are the following findings: In the UL field of older adults, Purdue pegboard accuracy (number of dropped pins) shows strong (*r* > .5) negative correlations with pQSM values in Ls and L5a layers. More iron in the Ls and L5a in older adults was therefore associated with more dropped pins (i.e., worse accuracy). In the F field in older adults, tongue movement errors (as measured by TT) show strong (*r* > .5) negative correlations with pQSM values in L5b and L6. More iron in L5b and L6 in older adults was therefore associated with more movement errors (i.e., worse accuracy). Conversely, the 6MWT scores did not show any strong (*r* > .5) correlations with the LL field in older adults. Our data therefore indicate that increased iron in older adults may have a link to poorer accuracy in motor tasks, while the layer-specific differences between cortical fields need further investigation.

**Figure 8.**
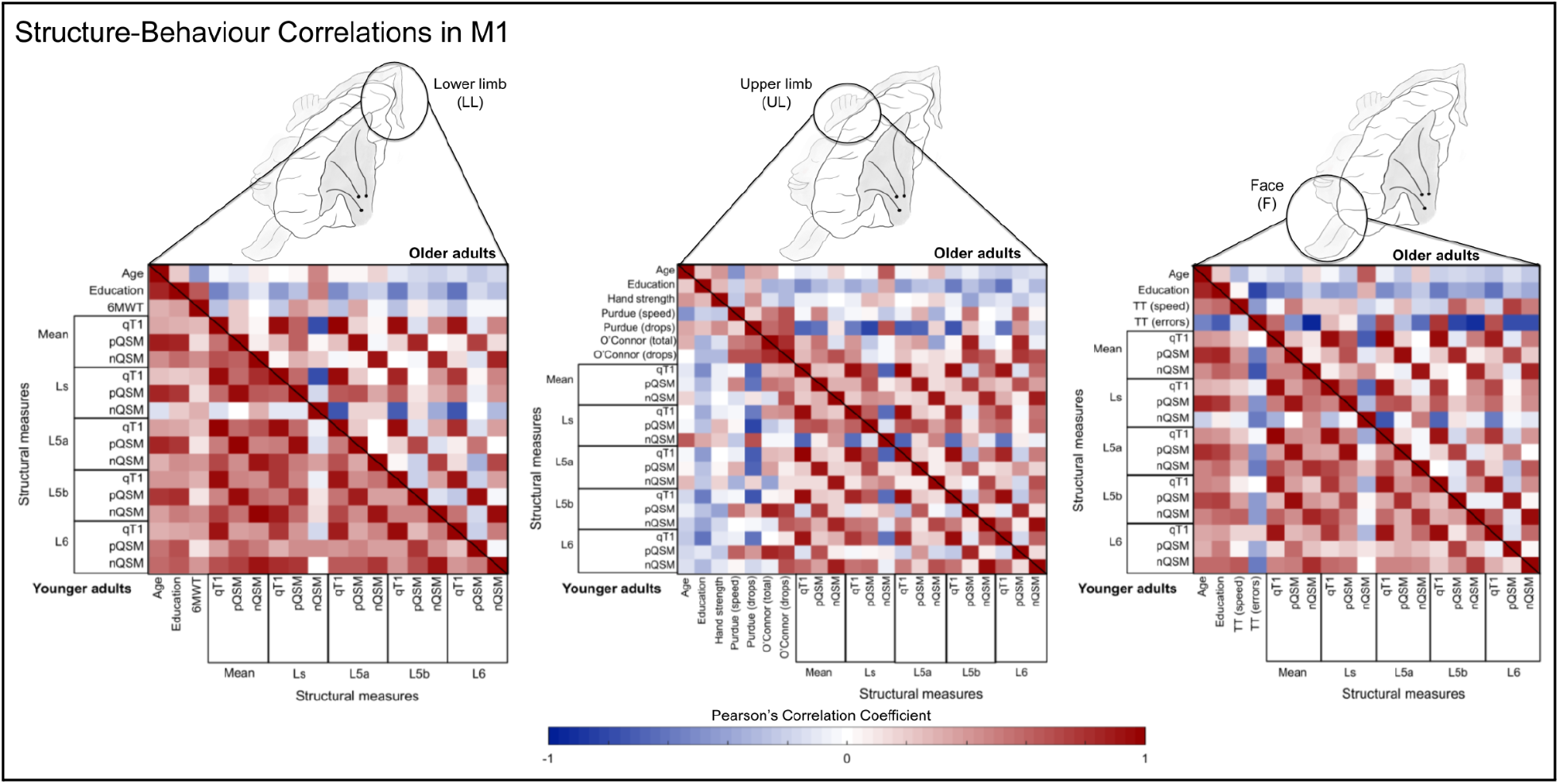
Correlations between M1 Microstructure and Body Part-Specific Motor Function. Correlation matrices between structural and behavioral measures are shown for each cortical field (lower limb, LL, upper limb, UL, face, F). The correlations for older adults are shown in the upper triangles of each matrix, while the correlations for younger adults are shown in the lower triangles of each matrix. Red colors indicate that ‘more substance’ relates to ‘better behavior’, whereas blue colors indicate that ‘more substance’ relates to ‘worse behavior’. For this purpose, Purdue pegboard scores (seconds to complete and dropped pins), O’Connor pegboard scores (dropped pins), tongue movement scores as estimated by the tongue tracker (TT) (duration and errors) and qT1 values were reversed, while nQSM values were taken as absolute values. The color bar shows the size of the correlation coefficient (Ls = superficial layer; L5a = layer 5a; L5b = layer 5b; L6 = layer 6).

## 4. Discussion

Topographic maps are a hallmark feature of the human brain, covering almost half of the cortical surface (Sereno et al., 2022). Cortical microcircuits of topographic maps, such as columns and layers, reveal critical insights into the mechanisms of aging and neurodegeneration, yet are poorly characterized in living older adults. We used M1 as a model system to characterize microstructural topographic map aging by applying parcellation-inspired techniques to sub-millimetre structural MRI data of healthy younger and older adults. We demonstrate systematic differences in the microstructural tissue profiles between the lower limb (LL), upper limb (UL) and face (F) areas in younger adults, suggesting that M1 is comprised of microstructurally distinct cortical fields. We show that these cortical fields are also distinct in older adults, and that the low-myelin borders between them do not degenerate in healthy aging. We also show that increased iron in older adults, a hallmark of cortical aging (Hallgren & Sourander, 1958), is particularly prominent in layer 5a of both hemispheres and in the LL field of left M1. Overall, we provide a novel 3D model of the microstructural architecture human M1, where layer-specific vulnerability is a central mechanism of cortical aging.

In the present study, we used a data-driven approach to estimate anatomically-relevant cortical compartments using structural data with a sub-millimeter isotropic resolution of 0.5 mm (explained in more detail in section 2.5.6). It must be noted that there is a conceptual difference between our definition of layers and the definition based on *ex-vivo* data, where cytoarchitecture is considered (Brodmann, 1909; Vogt & Vogt, 1919). Nevertheless, our approach is based on a comparison between *in-vivo* and *ex-vivo* M1 data (Huber et al., 2017) and we were able to characterize intracortical contrast with high precision (see Fig. 2). We therefore provide a reasonable approximation of layers and their computations (Persichetti et al., 2020). While previous studies often distinguish between input and output structures, we extended this by distinguishing between layer 5a and layer 5b. We are cautious in our interpretation of the different effects within layer 5, however, and therefore draw conclusions more generally about the output layer 5. Future studies should apply this layering approach to *ex-vivo* data, where also cytoarchitectural data exist, to test the reliability of this measure.

We here provide a novel model of the human 3D microstructure in M1 (see **Fig. 4A**). More precisely, we show that quantitative markers of cortical microstructure significantly differ between body parts, which we interpret to show that M1 is comprised of microstructurally distinct cortical fields that represent major body parts. These differences sometimes occur across cortical layers (e.g. the F field is characterized by high iron content and low nQSM) while they are sometimes layer-specific (e.g. the F field shows low myelin in L6 compared to the UL and LL fields). Our findings challenge classical depictions of human M1 as a single cortical field (Brodmann, 1909), suggesting instead that M1 is comprised of distinct cortical fields, each with distinct microstructural and functional features (Flechsig, 1920; Sereno et al., 2022). These microstructural differences may relate to differences in the cytoarchitecture and associated myeloarchitecture between cortical fields, for example where the size, shape and clustering of the heavily-myelinated Betz cells differ along the M1 strip (Rivara et al., 2003), and where differences in the myeloarchitectonic patterns have been identified (Flechsig, 1920).

Critically, we do not find an interaction between age and cortical field in myelin (qT1) or diamagnetic substance (nQSM) content, suggesting that the differences between cortical fields are also present in older adults. In addition, we do not show evidence of degenerated low-myelin borders between the hand and the face area in M1 in older adults. Our hypothesis that a degenerated myelin border relates to more functional overlap is therefore not confirmed by our data. The finding that cortical fields remain microstructurally distinct in healthy aging highlights limits to age-related plasticity within M1 (Sereno, 2005), providing a mechanistic explanation as to why maps are often maintained after deprivation (Makin & Bensmaia, 2017; Striem-Amit et al., 2018), and why body representations are often preserved in older age (Riemer et al., 2019). Moreover, since the low-myelin borders differ between individuals, they may explain the high inter-individual variability in neurodegenerative diseases involving topographic disease spread (Schreiber et al., 2021).

While previous studies have highlighted the vulnerability of M1 to age-related iron accumulation (Hallgren & Sourander, 1958; Acosta-Cabronero et al., 2016; Betts et al., 2016), we extend these findings to show that this occurs unevenly across cortical depth in M1. More specifically, we demonstrate that age-related increases in iron occur most strongly in what we defined as layer 5a of the left hemisphere. Layer 5, which is here divided into layer 5a and layer 5b, is responsible for the output of motor function via the heavily-myelinated Betz cells (McColgan et al., 2020). Interestingly, iron accumulation in the deeper L5b is often implicated in neurodegenerative diseases affecting motor control (McColgan et al., 2020). Our findings suggest that age-related differences in iron accumulation within the output layer 5 may distinguish healthy aging from neurodegeneration in M1. Further research, perhaps using invasive techniques in animal models, should investigate the effects of aging on layer 5 in more detail.

In addition to age-related increases in iron, we also demonstrate that the superficial layer and layer 5a of M1 show lower nQSM (that is, more substance) in older adults compared to younger adults. The biological source of nQSM is currently under debate but has been shown to reflect myelin (Deh et al., 2018) and calcium (Wang et al., 2017; Jang et al., 2021; Kim et al., 2022). Our data support the latter interpretation, by showing age-related increase in nQSM signal (i.e. more negative) without age-related differences in a validated proxy of myelin content (Stüber et al., 2014). Calcium deposits are well-evidenced in healthy aging and are associated with negative consequences on cognitive function (Thibault et al., 2007; Toescu & Verkhratsky, 2007). Interestingly, calcium deposits in older adults have been shown to occur largely near vascular structures (Jang et al., 2021), which may explain why the age-related increase in nQSM signal was strongest in the superficial layer of M1, where large vessels are located.

There is also a complex relationship between magnetic susceptibility and amyloid beta accumulation. While amyloid accumulation has been shown to be diamagnetic using *ex-vivo* 7T-MRI (Gong et al., 2019), it has also been shown to give rise to paramagnetic signal change based on a more specific approach comparing *ex-vivo* MRI at 9.4T and 14.1T (Tuzzi et al., 2020), where the latter may relate to iron accumulation near amyloid deposits. It is unclear whether our data support the interpretation of age-related nQSM increases as amyloid accumulation, given that healthy older adults would be expected to show minor or no amyloid accumulation in M1. Note that age-related and layer-specific differences cannot be due to group differences in layer width, since the size of the layer compartments was equal across age groups, even though they were defined separately within each group. These results are also unlikely to be explained by age-related differences in cortex morphology, since other microstructural tissue profiles that were extracted using the same masks, such as qT1, are stable across age groups. In addition, a control analysis showed that even when taking individual mask size into account, the age-related iron difference is still significant. Taken together, with strongest iron accumulation in layer 5, we here indicate that layer-specific vulnerability is a central mechanism of topographic map aging.

In terms of age-related differences in iron across cortical fields of M1, we show a small effect, where the LL field shows the highest effect and the UL field shows the smallest. This effect is only significant in the left (dominant) hemisphere. However, this finding is supported by animal research, where the hindpaw, but not the forepaw, representation of rats shows early signs of aging (David-Jürgens et al., 2008). This may explain why older adults show particularly deteriorated walking behavior, which is associated with reduced independence in late life (de Bruin & Schmidt, 2010). Since the UL field shows the smallest age effect in the dominant hemisphere, we suggest that use-dependent plasticity (i.e. of the dominant hand) may preserve topographic maps in the face of cortical aging. This is supported by the absence of differences in age-related iron accumulation between cortical fields of the right hemisphere (i.e. non-dominant body representations). This finding has some implications for basic research on the motor system, for example, it challenges the use of the hand area as a model system to study motor learning (Dumel et al., 2018) given the foot area of the dominant hemisphere may degenerate earlier. Our results are also relevant for neurodegenerative diseases where iron accumulation in specific cortical fields is considered as a disease-specific marker (Ravits & La Spada, 2009), and has been linked to symptom severity (Costagli et al., 2016; Kwan et al., 2012) and topographical disease spread (Schreiber et al., 2021). The present study provides a comprehensive model of the 3D microstructure of M1 in younger and older adults, to be used as a reference point for defining disease-specific features of pathology.

Finally, we performed correlational analyses to investigate how (age-related differences in) the cortical microstructure of M1 relates to motor function. It is worth noting that these analyses were done on reduced sample sizes for which behavioral data were available (n = 12 for younger adults, n = 15 for older adults). These preliminary analyses indicate that more iron in older adults relates to worse behavioral accuracy, and that the relationships between iron and motor behavior may be layer-dependent in older adults. More specifically, we show that increased iron in the superficial layer and layer 5a of the UL field is strongly associated with decreased accuracy in hand movements (*r* > .5), whereas increased iron in the layer 5b and layer 6 is associated with decreased accuracy in the tongue movement task (*r* > .5). This is in line with previous evidence of reduced hand dexterity with iron accumulation in aging (Li et al., 2015). Why in different cortical fields, more iron in specific layers relates to decreased accuracy needs to be clarified by future research. In both age groups, we found no strong correlations (*r* > .5) between microstructural measures and performance on the 6MWT, suggesting that this measure is not specific enough to relate to microstructural measures of the LL field. This may be due to the fact that we extracted microstructural profiles from each hemisphere separately, but walking is a bilateral motor task involving different body parts such as the foot, the limb and the knee, and where also the structure of the corpus callosum may be influential. Further research should use larger sample sizes to further describe the relationships between layer-specific iron content and motor function.

## 5. Conclusions

Taken together, our study highlights the importance of a 3D approach to topographic map aging that recognises layers and cortical fields as functionally-relevant units (Kuehn & Sereno, 2018), which show distinct age effects. Using recent advances in *in-vivo* microstructural imaging, we present a novel model of human M1 microstructure that includes distinct cortical fields and low-myelin borders, both in younger and older adults. We highlight that iron accumulation in layer 5, and diamagnetic substance (calcium) accumulation in layer 5 and the superficial layer, demonstrate layer-specific vulnerability in cortical aging. We suggest that further work should aim to disentangle the cortical field-specific iron accumulation in aging from that of neurodegeneration, where this is currently considered a disease-specific marker. We argue that layer-specific vulnerability is a central mechanism of topographic map aging, which inspires novel therapeutic interventions in M1 and beyond.

## Data Availability

Anonymised data can be made available upon request.

## Author Contributions

E.K., S.S. and M.W. designed the study. J.D., M.W. and A.N. collected the data. A.N. and J.D. completed the data processing. A.N. carried out the data investigation and statistical analysis. A.N. wrote the manuscript. J.D., E.K., S.S and M.W. reviewed and edited the manuscript. E.K., S.S. and S.V. supervised the study. All authors contributed to the article and approved the submitted version.

## Acknowledgements

We would like to thank Lilith-Sophie Lange for her support in data collection. This project was funded by the Else Kröner Fresenius Stiftung: 2019-A03 and the Deutsche Forschungsgemeinschaft (DFG): KU 3711/2-1 (project number 423633679), SCHR 1418/5-1 (501214112) and work package B04 (425899996) in CRC 1436.

