## Supplementary materials for "Layer-Specific Vulnerability is a Mechanism of Topographic Map Aging"

| | Young | Old | | | | |
|---|---|---|---|---|---|---|
| | **M ± SD** | **M ± SD** | ***df*** | ***t*** | ***Sig.*** | ***g, [95% CI]*** |
| 6MWT | 784.52 ± 53.15 | 449.06 ± 46.61 | 20 | 15.78 | $10^{-13}$* | 6.50 [4.33 8.64] |
| Hand strength (kg) | 37.50 ± 7.61 | 27.01 ± 9.02 | 26 | 3.30 | .003* | 1.21 [.41 2.00] |
| Purdue (secs) | 57.50 ± 7.25 | 62.90 ± 10.08 | 18 | -1.40 | .178 | -.60 [-1.45 .27] |
| Purdue (drops) | .70 ± .68 | .80 ± .63 | 18 | -.34 | .736 | -.15 [-.99 .70] |
| O’Connor (total) | 36.80 ± 7.76 | 32.70 ± 9.46 | 18 | 1.06 | .303 | .45 [-.41 1.30] |
| O’Connor (drops) | 7.60 ± 4.20 | 4.40 ± 3.81 | 18 | 1.79 | .091 | .77 [-.12 1.63] |
| TT (speed) | .045 ± .079 | .063 ± .061 | 17 | -.56 | .584 | -.25 [-1.11 .62] |
| TT (errors) | 6.40 ± 9.05 | 2.22 ± 3.03 | 17 | 1.32 | .205 | .58 [-.31 1.45] |

**Supplementary Table 1.** Group differences in behavioral tests of motor function. We quantified motor function of the legs (6MWT), hands (hand strength, Purdue, O’Connor) and face/tongue (TT). 6MWT = 6-Minute Walk Test. Purdue = Purdue pegboard test. O’Connor = O’Connor pegboard test. TT = Tongue Tracker. * indicates statistical significance at 5% level, corrected for multiple comparisons using the Holm-Bonferroni method.

| Factor | *df* | Sum of squares | Mean square | *F* | *p* | $\eta_p^2$ |
|---|---|---|---|---|---|---|
| Layer | 1.39 | $10^{-6}$ | $10^{-6}$ | 850.41 | $10^{-21}$** | .98 |
| Cortical field | 2 | 974.50 | 487.25 | .97 | .392 | .06 |
| Layer * cortical field | 2.30 | 2659.76 | 1157.54 | 7.20 | .002 | .31 |

**Supplementary Table 2.** Main effects of layer and cortical field, and the interaction between layer and cortical field on qT1 values in the right hemisphere of younger (n = 17) adults. * and ** indicate significance at the 5% and 1% significance levels, respectively. Degrees of freedom reported as non-integers indicate that equal variances were not assumed based on Levene’s test for equality of variances.

| Pair | Paired differences | | | | | |
| --- | --- | --- | --- | --- | --- | --- |
| | ***df*** | **Mean difference** | **Standard deviation** | ***t*** | ***Sig.*** | ***d, [95% CI]*** |
| Layer | | | | | | |
| Ls-L5a | 16 | 191.91 | 44.03 | 17.97 | $10^{-12}$* | 4.36 [2.78 5.92] |
| L5a-L5b | 16 | 156.85 | 23.61 | 27.39 | $10^{-15}$* | 6.64 [4.31 8.97] |
| L5b-L6 | 16 | 169.75 | 24.50 | 28.57 | $10^{-15}$* | 6.93 [4.50 9.35] |
| Layer * cortical field | | | | | | |
| Ls: F-UL | 16 | 1.54 | 25.70 | .25 | .808 | .06 [-.42 .54] |
| Ls: UL-LL | 16 | -9.82 | 21.69 | -1.87 | .080 | -.45 [-.95 .05] |
| Ls: LL-F | 16 | 8.27 | 19.65 | 1.74 | .102 | .42 [-.08 .91] |
| L5a: F-UL | 16 | -6.59 | 20.41 | -1.33 | .202 | -.32 [-.81 .17] |
| L5a: UL-LL | 16 | -.69 | 13.04 | -.22 | .829 | -.05 [-.53 .42] |
| L5a: LL-F | 16 | 7.28 | 17.93 | 1.67 | .114 | .41 [-.10 .90] |
| L5b: F-UL | 16 | -4.58 | 18.48 | -1.02 | .332 | -.25 [-.73 .24] |
| L5b: UL-LL | 16 | -1.65 | 12.85 | -.53 | .603 | -.13 [-.60 .35] |
| L5b: LL-F | 16 | 6.23 | 18.62 | 1.38 | .187 | .34 [-.16 .82] |
| L6: F-UL | 16 | 14.43 | 19.57 | 3.04 | .008 | .74 [.19 1.27] |
| L6: UL-LL | 16 | -8.31 | 13.10 | -2.62 | .019 | -.64 [-1.15 -.10] |
| L6: LL-F | 16 | -6.12 | 17.46 | -1.45 | .168 | -.35 [-.84 .15] |

**Supplementary Table 3.** Post-hoc t-tests for significant ANOVA effects on qT1 values in the right hemisphere of younger (n = 17) adults. (Ls = superficial layer; L5a = layer 5a; L5b = layer 5b; L6 = layer 6). * indicates statistical significance at 5% level, corrected for multiple comparisons using the Holm-Bonferroni method.

| Factor | *df* | Sum of squares | Mean square | *F* | *p* | $\eta_p^2$ |
|---|---|---|---|---|---|---|
| Layer | 1.25 | .002 | .002 | 178.15 | $10^{-10}$** | .93 |
| Cortical field | 2 | $10^{-5}$ | $10^{-6}$ | 11.45 | $10^{-4}$** | .47 |
| Layer * cortical field | 2.54 | $10^{-6}$ | $10^{-7}$ | 7.30 | $10^{-3}$** | .36 |

**Supplementary Table 4.** Main effects of layer and cortical field, and the interaction between layer and cortical field on pQSM values in the right hemisphere of younger (n = 14) adults. * and ** indicate significance at the 5% and 1% significance levels, respectively. Degrees of freedom reported as non-integers indicate that equal variances were not assumed based on Levene's test for equality of variances.

| Pair | Paired differences | | | | | |
|---|---|---|---|---|---|---|
| | ***df*** | **Mean difference** | **Standard deviation** | ***t*** | ***Sig.*** | ***d, [95% CI]*** |
| Layer | | | | | | |
| Ls-L5a | 13 | .002 | .0011 | 7.11 | $10^{-6}$* | 1.90 [1.00 2.78] |
| L5a-L5b | 13 | .004 | .0012 | 13.81 | $10^{-9}$* | 3.69 [2.19 5.18] |
| L5b-L6 | 13 | .002 | .0007 | 12.82 | $10^{-9}$* | 3.43 [2.02 4.82] |
| Layer * cortical field | | | | | | |
| Ls: F-UL | 13 | .0013 | .0009 | 5.04 | $10^{-4}$* | 1.35 [.60 2.07] |
| Ls: UL-LL | 13 | -.0006 | .0008 | -2.49 | .027* | -.67 [-1.24 -.07] |
| Ls: LL-F | 13 | -.0007 | .0008 | -3.23 | .007* | -.86 [-1.47 -.23] |
| L5s: F-UL | 13 | .0010 | .0008 | 4.55 | $10^{-4}$* | 1.22 [.51 1.90] |
| L5s: UL-LL | 13 | -.0004 | .0007 | -2.28 | .040* | -.61 [-1.17 -.03] |
| L5s: LL-F | 13 | -.0006 | .0007 | -2.79 | .015* | -.75 [-1.33 -.14] |
| L5b: F-UL | 13 | .0004 | .0006 | 2.24 | .043* | .60 [.02 1.16] |
| L5b: UL-LL | 13 | -.0002 | .0004 | -1.92 | .078 | -.51 [-1.06 .06] |
| L5b: LL-F | 13 | -.0002 | .0005 | -1.44 | .174 | -.38 [-.92 .17] |
| L6: F-UL | 13 | .0002 | .0007 | 1.10 | .290 | .30 [-.25 .83] |
| L6: UL-LL | 13 | -.0001 | .0006 | -.90 | .383 | -.24 [-.77 .30] |
| L6: LL-F | 13 | -.0001 | .0005 | -.48 | .641 | -.13 [-.65 .40] |

**Supplementary Table 5.** Post-hoc t-tests for significant ANOVA effects on pQSM values in the right hemisphere of younger (n = 14) adults. (Ls = superficial layer; L5a = layer 5a; L5b = layer 5b; L6 = layer 6). * indicates statistical significance at 5% level, corrected for multiple comparisons using the Holm-Bonferroni method.

| Factor | *df* | Sum of squares | Mean square | *F* | *p* | $\eta_p^2$ |
|---|---|---|---|---|---|---|
| Layer | 1.16 | .001 | .001 | 83.20 | $10^{-8}$** | .87 |
| Cortical field | 2 | $10^{-6}$ | $10^{-6}$ | 3.68 | .039* | .22 |
| Layer * cortical field | 2.75 | $10^{-6}$ | $10^{-6}$ | 3.53 | .028* | .21 |

**Supplementary Table 6.** Main effects of layer and cortical field, and the interaction between layer and cortical field on nQSM values in the right hemisphere of younger (n = 14) adults. * and ** indicate significance at the 5% and 1% significance levels, respectively. Degrees of freedom reported as non-integers indicate that equal variances were not assumed based on Levene's test for equality of variances.

| Pair | Paired differences | | | | | |
|---|---|---|---|---|---|---|
| | ***df*** | **Mean difference** | **Standard deviation** | ***t*** | ***Sig.*** | ***d , [95% CI]*** |
| Layer; cortical field | | | | | | |
| Ls-L5a | 13 | -.0011 | .0014 | -3.02 | .010 | -.81 [-1.40 .19] |
| L5a-L5b | 13 | .0026 | .0010 | 9.55 | $10^{-7}$* | 2.55 [1.44 3.62] |
| L5b-L6 | 13 | .0043 | .0008 | 21.21 | $10^{-11}$* | 5.67 [3.45 7.88] |
| F-UL | 13 | -.0004 | .0007 | -2.36 | .034 | -.05 [-.57 .48] |
| UL-LL | 13 | $-10^{-5}$ | .0006 | -.19 | .856 | -.63 [-1.20 .05] |
| LL-F | 13 | .0004 | .0008 | 2.15 | .051 | .57 [$10^{-3}$ 1.07] |
| Layer * cortical field | | | | | | |
| Ls: F-UL | 13 | -.0010 | .0010 | -3.60 | .003* | -.96 [-1.59 .31] |
| Ls: UL-LL | 13 | .0004 | .0010 | 1.62 | .129 | .43 [-.12 .98] |
| Ls: LL-F | 13 | .0005 | .0011 | 1.67 | .119 | .45 [-.11 .99] |
| L5a: F-UL | 13 | -.0004 | .0005 | -2.83 | .014 | -.76 [-1.34 .15] |
| L5a: UL-LL | 13 | $-10^{-5}$ | .0007 | -.08 | .934 | -.02 [-.55 .50] |
| L5a: LL-F | 13 | .0004 | .0007 | 2.09 | .057 | .56 [-.02 1.12] |
| L5b: F-UL | 13 | $-10^{-5}$ | .0007 | -.48 | .643 | -.13 [-.65 .40] |
| L5b: UL-LL | 13 | -.0003 | .0007 | -1.65 | .122 | -.44 [-.99 .12] |
| L5b: LL-F | 13 | .0004 | .0007 | 2.03 | .063 | .54 [-.03 1.10] |
| L6: F-UL | 13 | -.0002 | .0007 | -.76 | .463 | -.20 [-.73 .33] |
| L6: UL-LL | 13 | -.0002 | .0008 | -1.19 | .255 | -.32 [-.85 .23] |
| L6: LL-F | 13 | .0005 | .0010 | 1.78 | .099 | .48 [-.09 1.02] |

**Supplementary Table 7.** Post-hoc tests for significant effects of layer and cortical field on nQSM values in the right hemisphere of younger (n = 14) adults. (Ls = superficial layer; L5a = layer 5a; L5b = layer 5b; L6 = layer 6). * indicates statistical significance at 5% level, corrected for multiple comparisons using the Holm-Bonferroni method.

| Factor | *df* | Sum of squares | Mean square | *F* | *p* | $\eta_p^2$ |
|---|---|---|---|---|---|---|
| Layer | 1.21 | $10^{-7}$ | $10^{-7}$ | 731.93 | $10^{-28}$** | .96 |
| Cortical field | 2 | 1316.47 | 658.23 | 1.70 | .191 | .05 |
| Group | 1 | $10^{-5}$ | $10^{-5}$ | 1.90 | .178 | .05 |
| Layer * cortical field | 3 | 5501.36 | 1835.02 | 16.65 | $10^{-9}$** | .34 |
| Layer * group | 1.21 | 55528.36 | 45871.32 | 2.46 | .119 | .07 |
| Cortical field * group | 2 | 1096.14 | 548.07 | 1.41 | .250 | .04 |
| Layer * cortical field * group | 3 | 766.55 | 255.69 | 2.32 | .080 | .07 |

**Supplementary Table 8.** Main effects of layer, cortical field and age and the interactions between layer, cortical field and age on qT1 values in the right hemisphere of younger (n = 17) and older (n = 18) adults. * and ** indicate significance at the 5% and 1% significance levels, respectively. Degrees of freedom reported as non-integers indicate that equal variances were not assumed based on Levene's test for equality of variances.

| Pair | Paired differences | | | | | |
|---|---|---|---|---|---|---|
| | ***df*** | **Mean difference** | **Standard deviation** | ***t*** | ***Sig.*** | ***d, [95% CI]*** |
| Layer | | | | | | |
| Ls-L5a | 34 | 218.95 | 80.50 | 16.09 | $10^{-17}$* | 2.72 [1.99 3.44] |
| L5a-L5b | 34 | 160.38 | 30.11 | 31.51 | $10^{-27}$* | 5.33 [4.02 6.63] |
| L5b-L6 | 34 | 157.56 | 27.82 | 33.50 | $10^{-27}$* | 5.66 [4.28 7.04] |
| Layer * cortical field | | | | | | |
| Ls: F-UL | 34 | -7.79 | 23.49 | -1.96 | .058 | -.33 [-.67 .01] |
| Ls: UL-LL | 34 | -2.75 | 20.12 | -.81 | .425 | -.14 [-.47 .20] |
| Ls: LL-F | 34 | 10.53 | 20.10 | 3.10 | .004* | .52 [.17 .87] |
| L5a: F-UL | 34 | -9.08 | 16.17 | -2.60 | .002* | -.56 [-.92 -.20] |
| L5a: UL-LL | 34 | 2.04 | 12.65 | .95 | .348 | .16 [-.17 .49] |
| L5a: LL-F | 34 | 7.04 | 16.04 | 2.60 | .014 | .44 [.09 .78] |
| L5b: F-UL | 34 | -5.22 | 15.67 | -1.97 | .057 | -.33 [-.67 .01] |
| L5b: UL-LL | 34 | -.09 | 11.28 | -.05 | .961 | -.01 [-.34 .32] |
| L5b: LL-F | 34 | 5.32 | 16.17 | 1.95 | .060 | .33 [-.01 .67] |
| L6: F-UL | 34 | 12.02 | 17.60 | 4.04 | $10^{-4}$* | .68 [.31 1.05] |
| L6: UL-LL | 34 | -6.49 | 12.18 | -3.15 | .003* | -.53 [-.88 -.18] |
| L6: LL-F | 34 | -5.53 | 15.86 | -2.06 | .047 | -.35 [-.69 -.01] |

**Supplementary Table 9.** Post-hoc tests for significant effects of layer and cortical field on qT1 values in the right hemisphere of younger (n = 17) and older (n = 18) adults. (Ls = superficial layer; L5a = layer 5a; L5b = layer 5b; L6 = layer 6). * indicates statistical significance at 5% level, corrected for multiple comparisons using the Holm-Bonferroni method.

| Factor | *df* | Sum of squares | Mean square | *F* | *p* | $\eta_p^2$ |
|---|---|---|---|---|---|---|
| Layer | 1.14 | .002 | .002 | 93.81 | $10^{-11}$** | .77 |
| Cortical field | 2 | $10^{-6}$ | $10^{-6}$ | 3.32 | .043* | .11 |
| Group | 1 | $10^{-5}$ | $10^{-5}$ | 3.95 | .057 | .12 |
| Layer * cortical field | 2.84 | $10^{-6}$ | $10^{-6}$ | 3.72 | .016* | .12 |
| Layer * group | 1.14 | $10^{-6}$ | $10^{-6}$ | .14 | .746 | .01 |
| Group * cortical field | 2 | $10^{-6}$ | $10^{-6}$ | 1.42 | .249 | .05 |
| Layer * cortical field * group | 2.84 | $10^{-6}$ | $10^{-7}$ | .68 | .562 | .02 |

**Supplementary Table 10.** Main effects of layer, cortical field and age and the interactions between layer, cortical field and age on nQSM values in the right hemisphere of younger (n = 14) and older (n = 15) adults. * and ** indicate significance at the 5% and 1% significance levels, respectively. Degrees of freedom reported as non-integers indicate that equal variances were not assumed based on Levene's test for equality of variances.

| Pair | Paired differences | | | | | |
|---|---|---|---|---|---|---|
| | *df* | Mean difference | Standard deviation | *t* | *Sig.* | *d, [95% CI]* |
| Layer; cortical field | | | | | | |
| Ls-L5a | 16 | -.00117 | .00201 | -3.20 | .003* | -.58 [-.97 -.19] |
| L5a-L5b | 16 | .00249 | .00135 | 10.10 | $10^{-11}$* | 1.85 [1.25 2.43] |
| L5b-L6 | 29 | .00418 | .00078 | 29.24 | $10^{-23}$* | 5.34 [6.75 3.92] |
| F-UL | 13 | -.00017 | .00061 | -1.56 | .129 | -.29 [-.65 .08] |
| UL-LL | 13 | -.00016 | .00081 | -1.08 | .289 | -.20 [-.56 .17] |
| LL-F | 13 | .00033 | .00075 | 2.43 | .021 | .44 [.07 .82] |
| Layer * cortical field | | | | | | |
| Ls: F-UL | 29 | -.00066 | .00102 | -3.55 | .001* | -.65 [-1.04 -.25] |
| Ls: UL-LL | 29 | .00010 | .00098 | .56 | .580 | .10 [-.26 .46] |
| Ls: LL-F | 29 | .00056 | .00122 | 2.50 | .018 | .46 [.08 .83] |
| L5a: F-UL | 29 | -.00018 | .00056 | -1.71 | .097 | -.31 [-.68 .06] |
| L5a: UL-LL | 29 | -.00007 | .00091 | -.42 | .677 | -.08 [-.44 .28] |
| L5a: LL-F | 29 | .00025 | .00089 | 1.51 | .142 | .28 [-.09 .64] |
| L5b: F-UL | 29 | .00007 | .00070 | .54 | .595 | .10 [-.26 .46] |
| L5b: UL-LL | 29 | -.00027 | .00094 | -1.57 | .127 | -.29 [-.65 .08] |
| L5b: LL-F | 29 | .00020 | .00082 | 1.33 | .193 | .24 [-.12 .60] |
| L6: F-UL | 29 | .00007 | .00106 | .37 | .712 | .07 [-.29 .43] |
| L6: UL-LL | 29 | -.00040 | .00112 | -1.94 | .062 | -.35 [-.72 .02] |
| L6: LL-F | 29 | .00032 | .00105 | 1.70 | .100 | .31 [-.06 .67] |

**Supplementary Table 11.** Post-hoc t-tests for significant ANOVA effects on nQSM values in the right hemisphere of younger (n = 14) and older (n = 16) adults. (Ls = superficial layer; L5a = layer 5a; L5b = layer 5b; L6 = layer 6). * indicates statistical significance at 5% level, corrected for multiple comparisons using the Holm-Bonferroni method.

| | Young | Old | | | | |
|---|---|---|---|---|---|---|
| | **M ± SD** | **M ± SD** | ***df*** | ***t*** | ***Sig.*** | ***g, [95% CI]*** |
| Ls | -738.44 ± 203.73 | -808.05 ± 316.36 | 29.33 | .75 | .447 | .25 [-.41 .91] |
| L5a | -514.88 ± 236.56 | -603.65 ± 353.71 | 32 | .85 | .402 | .29 [-.38 .94] |
| L5b | -341.60 ± 104.65 | -396.61 ± 245.06 | 23.57 | .86 | .394 | .28 [-.38 .94] |
| L6 | -324.60 ± 97.17 | -316.43 ± 91.36 | 32 | -.25 | .802 | -.09 [-.74 .57] |

**Supplementary Table 12. Right hemisphere hand-face myelin border is stable in older adults.** We show no significant differences in the size of the right hemisphere hand-face myelin border ((hand+face/2)-border) between younger (n = 17) and older (n = 18) adults, at the uncorrected 5% significance level. (Ls - superficial layer, L5a - layer 5a, L5b - layer 5b, L6- layer 6). Degrees of freedom reported as non-integers indicate that equal variances were not assumed based on Levene's test for equality of variances.

| Factor | *df* | Sum of squares | Mean square | *F* | *p* | $\eta_p^2$ |
|---|---|---|---|---|---|---|
| Layer | 1.48 | .007 | .005 | 398.02 | $10^{-25}$** | .93 |
| Cortical field | 1.54 | $10^{-5}$ | $10^{-5}$ | 24.84 | $10^{-8}$** | .47 |
| Group | 1 | .002 | .002 | 31.11 | $10^{-6}$** | .53 |
| Layer * cortical field | 2.70 | $10^{-5}$ | $10^{-6}$ | 16.22 | $10^{-8}$** | .37 |
| Layer * group | 1.48 | $10^{-4}$ | $10^{-4}$ | 17.83 | $10^{-5}$** | .39 |
| Group * cortical field | 1.54 | $10^{-6}$ | $10^{-6}$ | 2.85 | .081 | .09 |
| Layer * cortical field * group | 2.70 | $10^{-6}$ | $10^{-7}$ | 1.33 | .272 | .05 |

**Supplementary Table 13.** Main effects of layer, cortical field and age and the interactions between layer, cortical field and age on pQSM values in the right hemisphere of younger (n = 14) and older (n = 15) adults. * and ** indicate significance at the 5% and 1% significance levels, respectively. Degrees of freedom reported as non-integers indicate that equal variances were not assumed based on Levene's test for equality of variances.

| Pair | Paired differences | | | | | |
|---|---|---|---|---|---|---|
| | *df* | Mean difference | Standard deviation | *t* | *Sig.* | *d, [95% CI]* |
| Layer; cortical field | | | | | | |
| Ls-L5a | 29 | .00276 | .00190 | 7.96 | $10^{-9}$* | 1.45 [.93 1.96] |
| L5a-L5b | 29 | .00566 | .00180 | 17.26 | $10^{-17}$* | 3.15 [2.26 4.03] |
| L5b-L6 | 29 | .00291 | .00100 | 15.96 | $10^{-16}$* | 2.92 [2.08 3.74] |
| F-UL | 29 | .00062 | .00103 | 4.68 | $10^{-6}$* | 1.02 [.57 1.45] |
| UL-LL | 29 | -.00105 | .00103 | -5.57 | $10^{-5}$* | -.85 [-1.27 -.43] |
| LL-F | 29 | -.00042 | .00067 | -3.48 | .002* | -.64 [-1.02 -.24] |
| Layer * cortical field | | | | | | |
| Ls: F-UL | 29 | .00176 | .00130 | 7.37 | $10^{-5}$* | 1.35 [.84 1.84] |
| Ls: UL-LL | 29 | -.00105 | .00119 | -4.83 | $10^{-8}$* | -.88 [-1.27 -.45] |
| Ls: LL-F | 29 | -.00070 | .00087 | -4.40 | $10^{-4}$* | -.80 [-1.21 -.39] |
| L5a: F-UL | 29 | .00138 | .00126 | 5.99 | $10^{-4}$* | 1.09 [.63 1.54] |
| L5a: UL-LL | 29 | -.00075 | .00093 | -4.40 | $10^{-6}$* | -.80 [-1.21 -.39] |
| L5a: LL-F | 29 | -.00063 | .00088 | -3.92 | $10^{-4}$* | -.72 [-1.11 -.31] |
| L5b: F-UL | 29 | .00073 | .00104 | 3.86 | .001* | .70 [.30 1.10] |
| L5b: UL-LL | 29 | -.00047 | .00074 | -3.45 | .002* | -.63 [-1.02 -.23] |
| L5b: LL-F | 29 | -.00026 | .00069 | -2.07 | .047 | -.38 [-.75 -.01] |
| L6: F-UL | 29 | .00032 | .00108 | 1.65 | .110 | .30 [-.07 .66] |
| L6: UL-LL | 29 | -.00022 | .00078 | -1.57 | .128 | -.29 [-.65 .08] |
| L6: LL-F | 29 | -.00010 | .00072 | -.77 | .450 | -.14 [-.50 .22] |

**Supplementary Table 14.** Post-hoc tests for significant effects of layer and cortical field on pQSM values in the right hemisphere of younger (n = 14) and older (n = 16) adults. (Ls = superficial layer; L5a = layer 5a; L5b = layer 5b; L6 = layer 6). * indicates statistical significance at 5% level, corrected for multiple comparisons using the Holm-Bonferroni method.

| | Young | Old | | | | |
|---|---|---|---|---|---|---|
| | M ± SD | M ± SD | *df* | *t* | *Sig.* | *g, [95% CI]* |
| Overall | .00964 ± .00258 | .01413 ± .00181 | 28 | -5.58 | $10^{-6}$* | -1.99 [-2.84 -1.11] |
| Ls | .01398 ± .00353 | .02089 ± .00262 | 23.76 | -6.02 | $10^{-6}$* | -2.19 [-3.08 -1.28] |
| L5a | .01188 ± .00307 | .01754 ± .00233 | 28 | -5.73 | $10^{-6}$* | -2.04 [-2.91 -1.15] |
| L5b | .00758 ± .00211 | .01069 ± .00200 | 28 | -4.14 | $10^{-4}$* | -1.47 [-2.26 -.67] |
| L6 | .00511 ± .00204 | .00740 ± .00209 | 28 | -3.03 | .005 | -1.08 [-1.82 -.32] |
| F | .01000 ± .00274 | .01473 ± .00210 | 24.23 | -5.26 | $10^{-6}$* | -1.91 [-2.75 -1.04] |
| UL | .00930 ± .00254 | .013387 ± .00158 | 28 | -5.36 | $10^{-5}$* | -1.91 [-2.75 -1.04] |
| LL | .00962 ± .00251 | .01427 ± .00195 | 24.46 | -5.60 | $10^{-5}$* | -2.03 [-2.89 -1.14] |

**Supplementary Table 15.** Post-hoc tests for significant effects of age on pQSM values in the right hemisphere of younger (n = 14) and older (n = 16) adults. (Ls = superficial layer; L5a = layer 5a; L5b = layer 5b; L6 = layer 6). * indicates statistical significance at 5% level, corrected for multiple comparisons using the Holm-Bonferroni method. Degrees of freedom reported as non-integers indicate that equal variances were not assumed based on Levene's test for equality of variances.

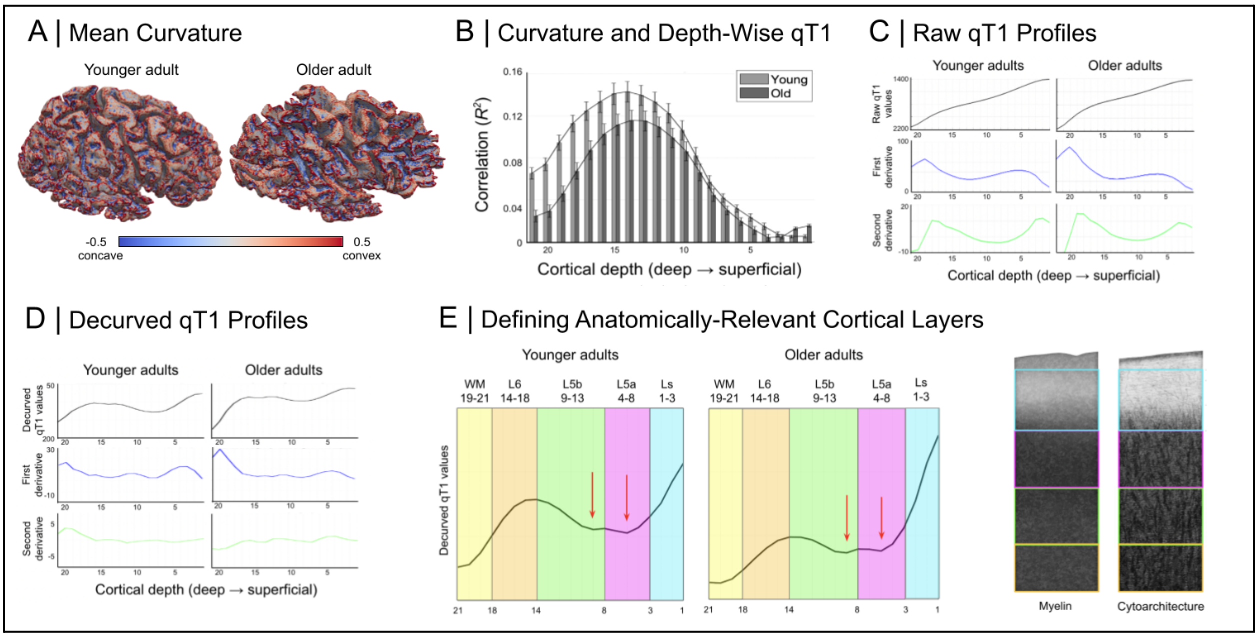


**Supplementary Figure 1. Identifying Anatomically-Relevant Cortical Compartments in Right Primary Motor Cortex (M1).** (A) Mapped mean cortical curvature for one example younger adult and one example older adult. (B) Cortical depth-wise correlations between qT1 values and mean curvature for younger adults (light gray) and older adults (dark gray). Note that the 21 correlations shown correspond to the initially-extracted 21 depths. (C) Group mean 'raw qT1' profiles along with the first and second derivatives. (D) Group mean 'decurved qT1' profiles along with the first and second derivatives. (E) According to a previously published approach (Huber et al., 2017), we identified four anatomically-relevant compartments ('layers') based on 'decurved qT1' (Ls - superficial layer; L5a - layer 5a; L5b - layer 5b; L6 - layer 6). L5a and L5b were distinguished based on the presence of two small qT1 dips at the plateau of 'decurved qT1' values (indicating L5), while L6 was identified based on a sharp decrease in values before a further plateau indicating the presence of white matter. We show our layer approximations over schematic depictions of M1 myelin (left) (from Dinse et al., 2015) and cell histological staining (right) (Vogt & Vogt, 1919). Note that these cortical layers are defined based on in-vivo MRI data and may not correspond exactly to the anatomical layers as defined by ex-vivo myelo- and cytoarchitecture. Also note that here, low qT1 values represent high myelin whereas in the other graphs in the article, values are plotted reversed such that high values represent high myelin.

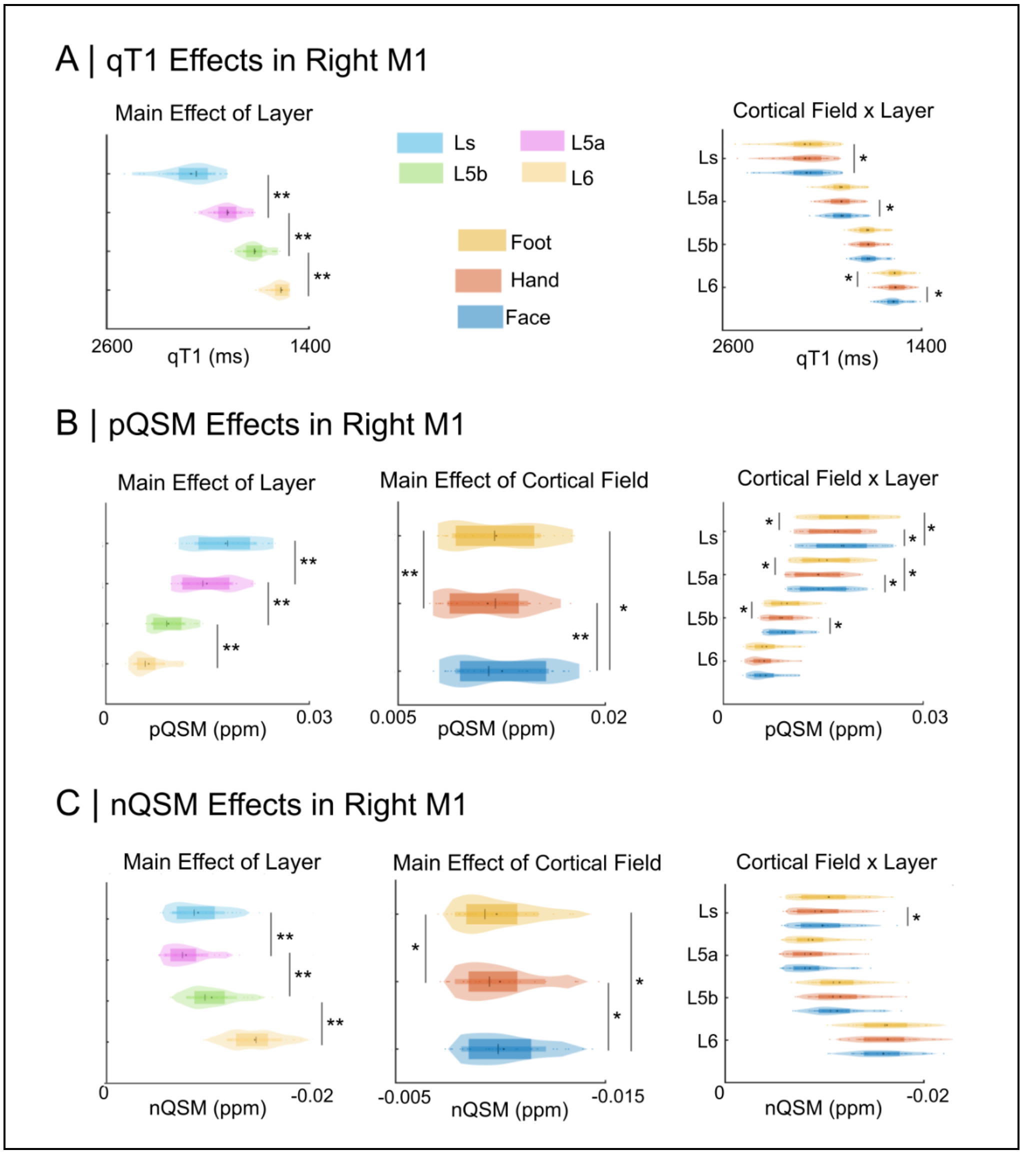


**Supplementary Figure 2. Microstructurally Distinct Cortical Fields in Right Primary Motor Cortex (M1)**. Layer- and cortical field-specific effects on (A) qT1 values as a proxy for cortical myelin content (Stuber et al., 2014), (B) pQSM values as a proxy of cortical iron content (Langkammer et al., 2012), and (C) nQSM values as a measure of diamagnetic tissue contrast (Deh et al., 2018). Main effects: * and ** indicate significance at the 5% and 1% significance levels, respectively. Interactions: * indicates statistical significance at 5% level, corrected for multiple comparisons using the Holm-Bonferroni method.

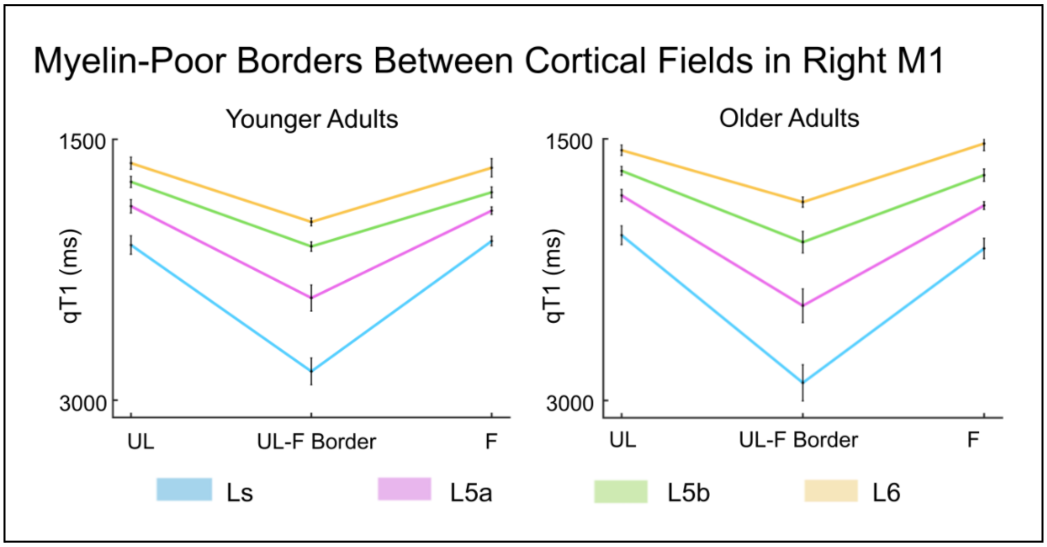


**Supplementary Figure 3.** Stable Hand-Face Myelin Borders in Older Adults in the Right Primary Motor Cortex (M1). Layer-wise low-myelin (high qT1 value) border between the cortical fields of the upper limb (UL) and face (F) for both younger and older adults. In each group, we compare the mean body part qT1 values (hand+face/2) with the qT1 values at the hand-face border across layers.

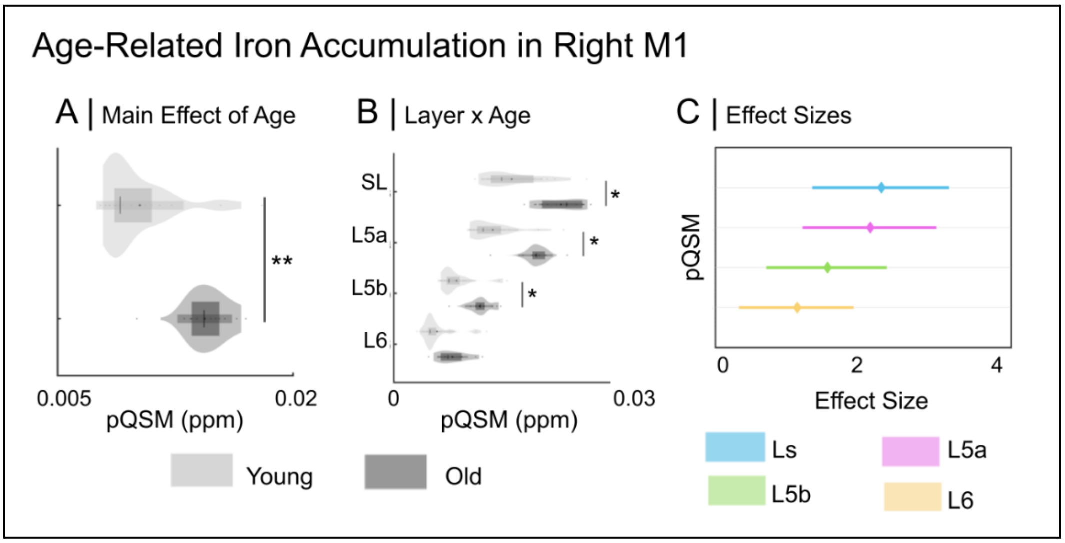


**Supplementary Figure 4. Age-Related Layer-Specific Iron Increases in Right Primary Motor Cortex (M1).** (A) Significant main effect of age and (B) significant interaction between layer and age on pQSM values (Ls = superficial layer; L5a = layer 5a; L5b = layer 5b; L6 = layer 6). (C) Effect sizes for the age effects on layer pQSM values. Main effects: * and ** indicate significance at the 5% and 1% significance levels, respectively. Interactions: * indicates statistical significance at 5% level, corrected for multiple comparisons using the Holm-Bonferroni method.
